# A rigorous benchmarking of alignment-based HLA typing algorithms for RNA-seq data

**DOI:** 10.1101/2023.05.22.541750

**Authors:** Dottie Yu, Ram Ayyala, Sarah Hany Sadek, Likhitha Chittampalli, Hafsa Farooq, Junghyun Jung, Abdullah Al Nahid, Grigore Boldirev, Mina Jung, Sungmin Park, Austin Nguyen, Alex Zelikovsky, Nicholas Mancuso, Jong Wha J. Joo, Reid F. Thompson, Houda Alachkar, Serghei Mangul

## Abstract

Accurate identification of human leukocyte antigen (HLA) alleles is essential for various clinical and research applications, such as transplant matching and drug sensitivities. Recent advances in RNA-seq technology have made it possible to impute HLA types from sequencing data, spurring the development of a large number of computational HLA typing tools. However, the relative performance of these tools is unknown, limiting the ability for clinical and biomedical research to make informed choices regarding which tools to use. Here we report the study design of a comprehensive benchmarking of the performance of 12 HLA callers across 682 RNA-seq samples from 8 datasets with molecularly defined gold standard at 5 loci, HLA-A, -B, -C, -DRB1, and -DQB1. For each HLA typing tool, we will comprehensively assess their accuracy, compare default with optimized parameters, and examine for discrepancies in accuracy at the allele and loci levels. We will also evaluate the computational expense of each HLA caller measured in terms of CPU time and RAM. We also plan to evaluate the influence of read length over the HLA region on accuracy for each tool. Most notably, we will examine the performance of HLA callers across European and African groups, to determine discrepancies in accuracy associated with ancestry. We hypothesize that RNA-Seq HLA callers are capable of returning high-quality results, but the tools that offer a good balance between accuracy and computational expensiveness for all ancestry groups are yet to be developed. We believe that our study will provide clinicians and researchers with clear guidance to inform their selection of an appropriate HLA caller.

## Introduction

The Human Leukocyte Antigen complex (HLA), otherwise known as the human Major Histocompatibility Complex (MHC), is a 3.6 Mb segment of the human genome which encodes for cell surface glycoproteins involved in cell-mediated immunity and self/nonself recognition. There are two classes of the HLA system typically implicated in immunity: Class I, which includes the loci HLA-A, HLA-B, and HLA-C and Class II, which includes the loci HLA-DRB1 and HLA-DQB1 (Fig 1). These five loci comprise the Classical HLA loci and are the most well documented within the entire region. The HLA region is the most polymorphic region of the human genome with over 25,000 HLA alleles currently available in the Immuno Polymorphism Database ImMunoGeneTics (IPD-IMGT) HLA reference database^1,2^. The high levels of polymorphism has the biological function of allowing each individual to produce a dramatically distinct immune response.

**Figure 1:**
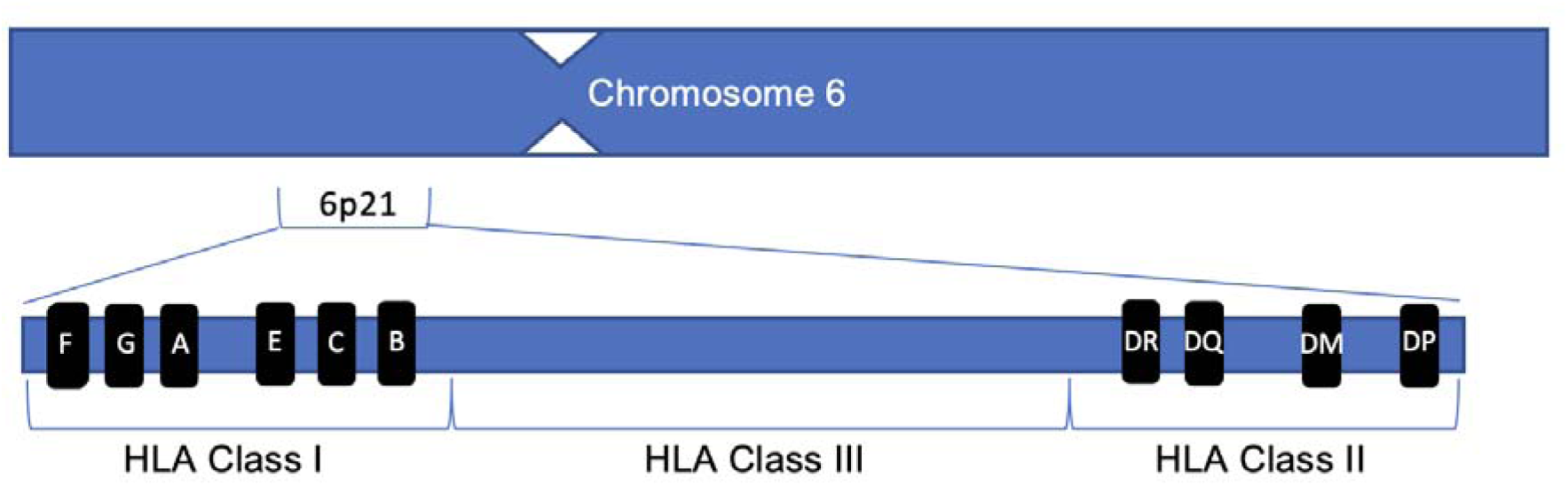
Schematic representation of HLA locus, with classical genes in Class I and Class II labeled.

**Figure 2:**
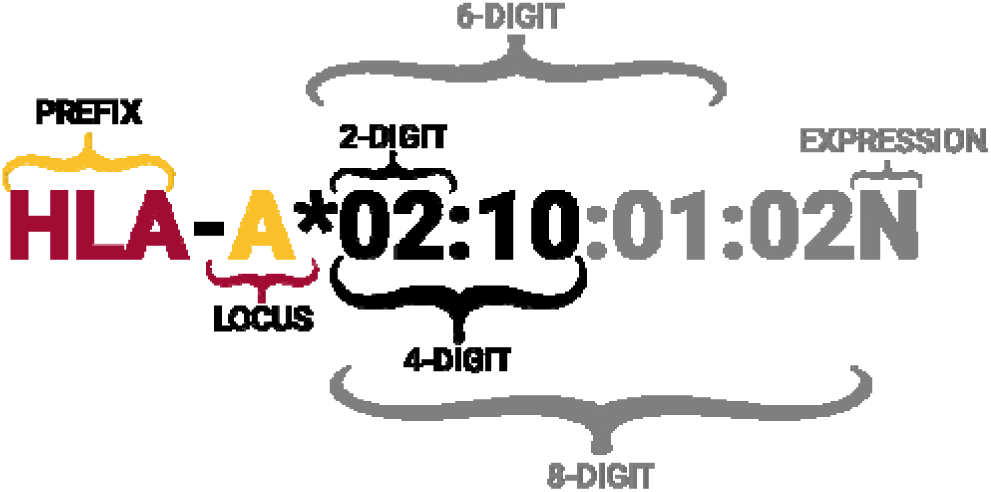
The nomenclature for specification of HLA allele types. One-field resolution indicates the allele group and is the lowest resolution for typing. Two-field resolution indicates the specific protein produced and is considered high resolution and the clinically significant resolution. Full resolution indicates synonymous DNA substitution in the coding region as well as changes in the non-coding areas of the genome.

HLA genes influence many immune-related disease phenotypes. Certain HLA variants and single nucleotide polymorphisms are associated with pathogenesis of both infectious and autoimmune diseases ^3,4^, adverse reactions to certain drugs^5^, and cancer development^6^. Recent studies have also found that some HLA variations are associated with susceptibility, progression, and severity of SARS-CoV-2 infection^7,8^. Therefore, accurate identification of HLA haplotypes is highly relevant to immune-related disease research. Furthermore, in the clinical setting, HLA typing is nevertheless widely used in transplant medicine including organ and hematopoietic stem cell transplantations^9^. However, it is worth noting that most RNA-seq based HLA typing methods are rarely used in the clinical setting, where the medical consequences of mistyping outweigh the advantages of NGS-based HLA typing. Nevertheless, US standards require an 8 out of 8 allele match for all haplotypes within the HLA-A, HLA-B, HLA-C, and HLA-DQB1 loci for transplant donor selection, and most European standards require a 10 out of 10 allele match for all haplotypes between the HLA-A, HLA-B, HLA-C, HLA-DRB1, and HLA-DQB1 loci. Whether a donor’s HLA type constitutes an “HLA match” depends on the resolution considered, with common resolutions being either one-field resolution (previously “two-digit”) or two-field resolution (previously “four-digit”) (Fig S1). High resolution typing is recommended by the National Marrow Donors Program and hence is of considerable clinical significance^10^.

Established HLA typing methods involve laboratory-based techniques such as amplification and quantification of HLA genes via sequence-specific oligonucleotide probes (SSOP) or sequence-specific primers (SSP), or via PCR followed by Sanger sequencing and comparison against the IPD-IMGT/HLA reference (SBT). SBT is one of the current gold standard methods of typing, both it is still cost prohibitive and laborious to be effective for large-scale clinical and research applications^7^. Recent developments in accessibility and throughput of NGS technology have spurred the development of HLA callers that impute HLA types from short read data. They use algorithms that can be divided into two general categories, alignment-based methods and assembly methods. Alignment-based methods align reads to an HLA reference containing a set of known allele sequences, then use that information to either build a probabilistic or optimization model to impute the HLA type. The less common assembly method used by HLAminer targeted assembly mode^11^, ATHLETES^12^, and HLAreporter^13^, first assembles sequencing reads into contigs which are then aligned to the reference genome to make the imputation of HLA type. Furthermore, another approach are the SNP-based HLA imputation methods SNP2HLA, HIBAG, HLA-check, and hlabud, which have been demonstrated to be useful for large-scale genetic studies on immune-disease related phenotypes^14,15^.

Despite the plethora of recently developed HLA callers, there is a lack of comprehensive and systematic benchmarking of RNA-seq-based HLA callers using large-scale and realist gold standards, limiting the ability for clinical and biomedical research to make informed choices regarding which tools to use. Furthermore, most existing benchmarking papers focus solely on whole-genome (WGS) or whole-exome (WES) data^16–20^, with the only existing study performed with RNA-seq samples completed in 2016^21^. Since that previous benchmarking study, seven new HLA callers have been developed, and the accuracy of HLA callers on varying read length and ancestry is yet to be evaluated. To address these limitations, we will evaluate the performance of 12 HLA callers currently available across 652 RNA-seq samples with available gold standard HLA alleles. The tools we plan to include in this analysis are HLAminer (read alignment mode)^11^, seq2HLA^22^, HLAforest^23^, PHLAT^24^, OptiType^25^, HLA-VBseq^26^, HLA-HD^27^, HISAT-genotype^28^, arcasHLA^29^, HLApers^30^, RNA2HLA^31^, T1K^32^, of which 8 are novel and not considered in the previous benchmarking study. All tools make Class I and Class II predictions with the exception of OptiType, which only makes Class I predictions. In each case, we plan to produce evaluation metrics of accuracy that is the percentage of correctly predicted alleles to determine the best performing tool. We will also conduct comparisons of the performance of each HLA caller across the various HLA genes to detect the genes that are consistently miscalled by HLA calling methods. Furthermore, we will examine the impact of sequencing parameters on the performance of each HLA caller and thoroughly assess the computational resources required for each software. Additionally, we will compare the performance of HLA methods across different ancestry groups and identify tools and trends in prediction accuracy and quality across samples of European and African ancestry. In summary, our aim is to conduct an extensive evaluation of all widely used alignment-based HLA callers, revealing valuable insights for researchers and clinicians to make an informed decision selecting the most appropriate RNA-Seq based HLA caller, offering a balance of consistency, performance, and scalability for their clinical and scientific needs.

## Results

We assembled 8 datasets comprising 682 bulk RNA-seq samples and 20 single cell RNA-seq samples (Table 1). The samples include lymphoblastoid cell lines (n=540), peripheral blood mononuclear cells (PBMCs) (N =112), bone marrow mononuclear cells (n=20), and whole blood (n=10). Samples have read lengths between 32bp and 126bp and raw read count between 29.9 million to 774 million.

**Table 1:**
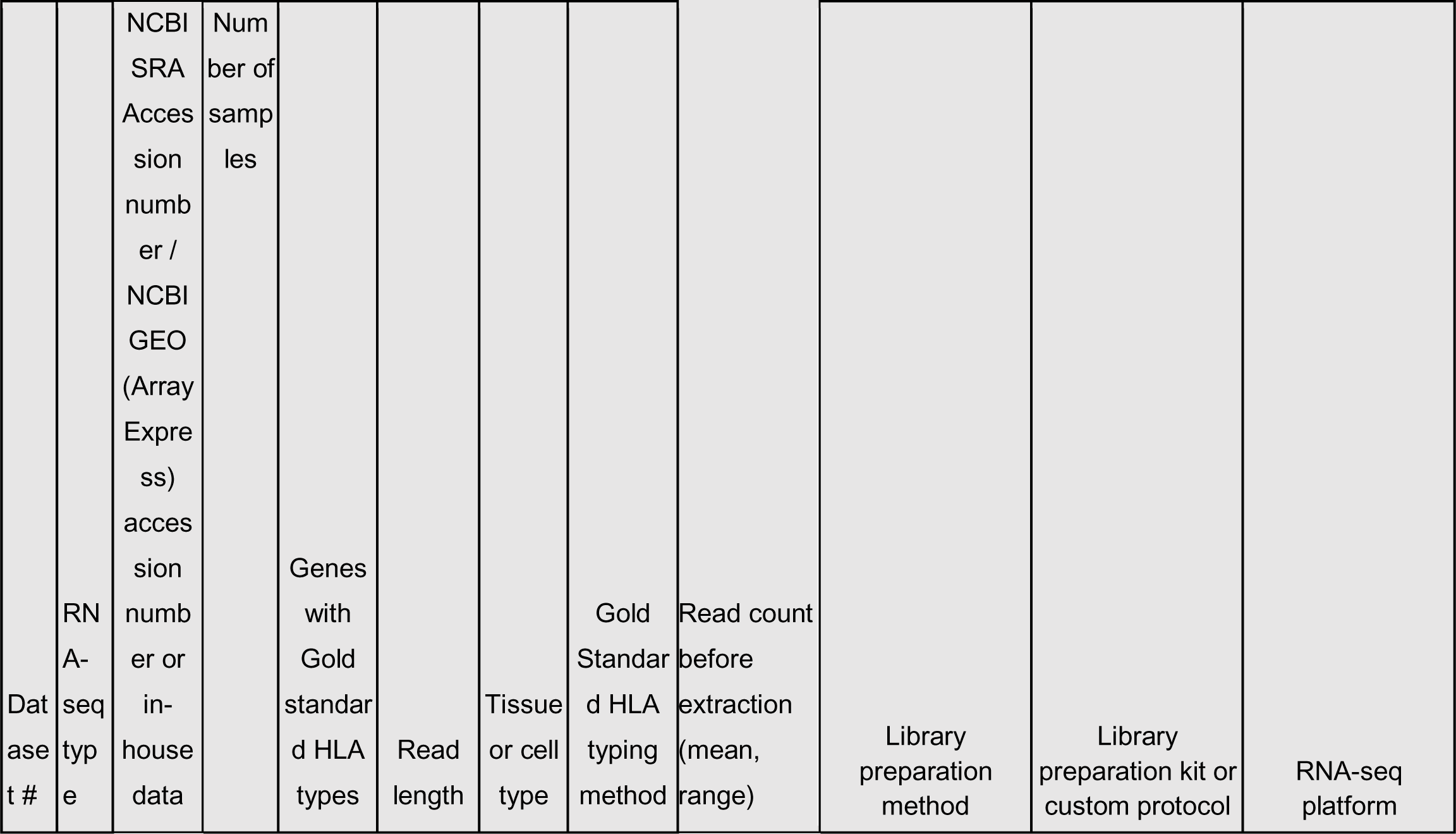
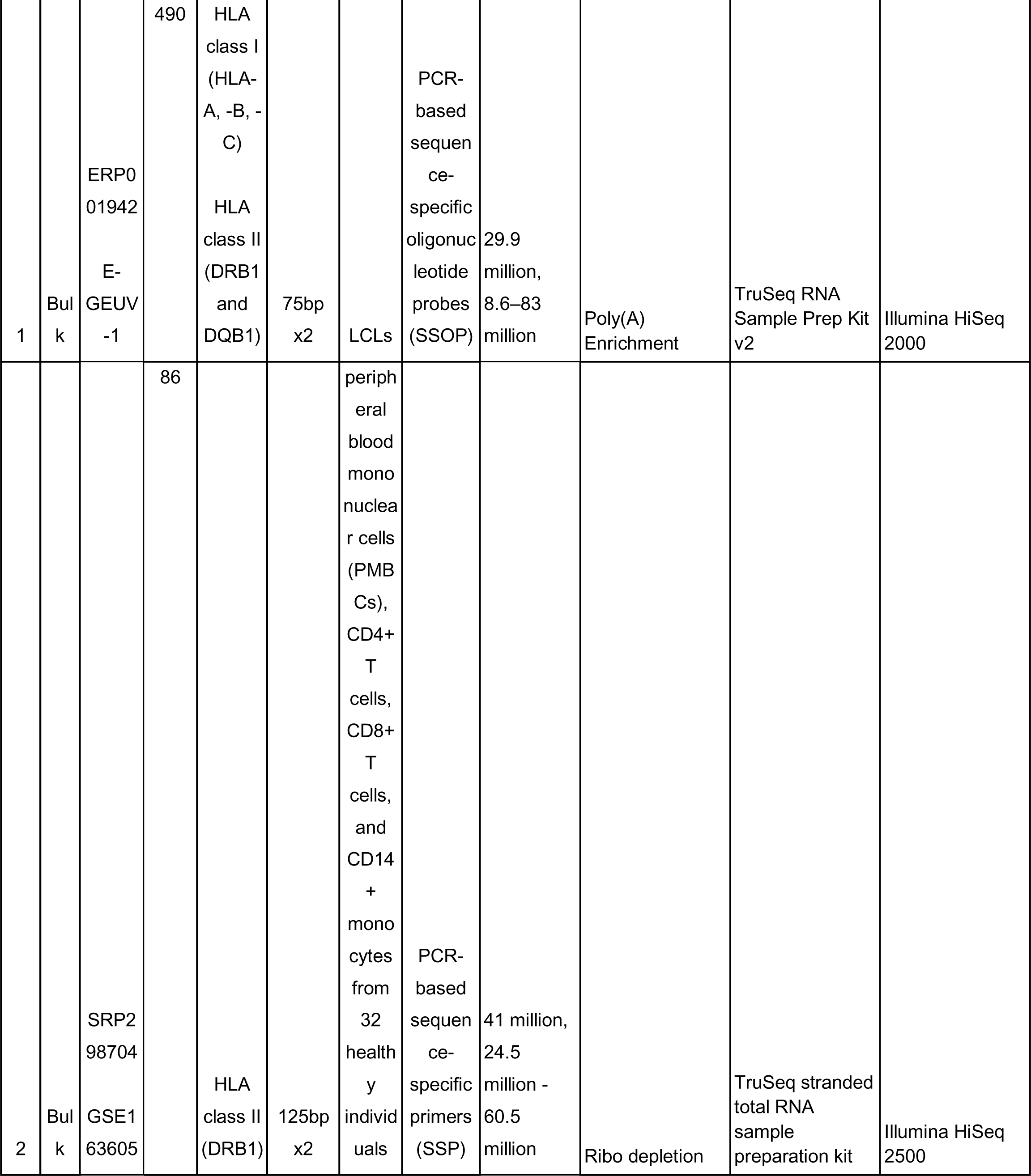
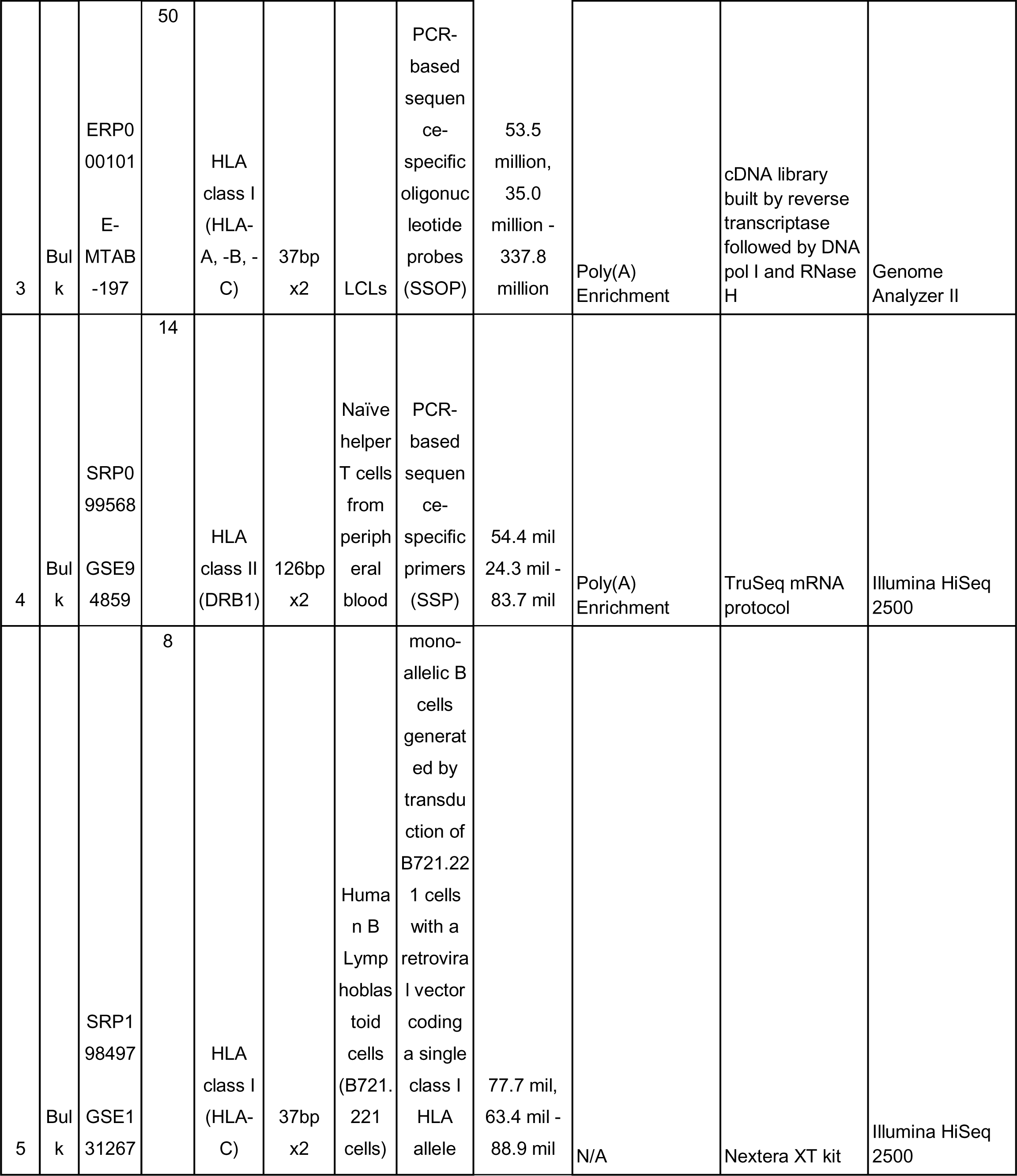
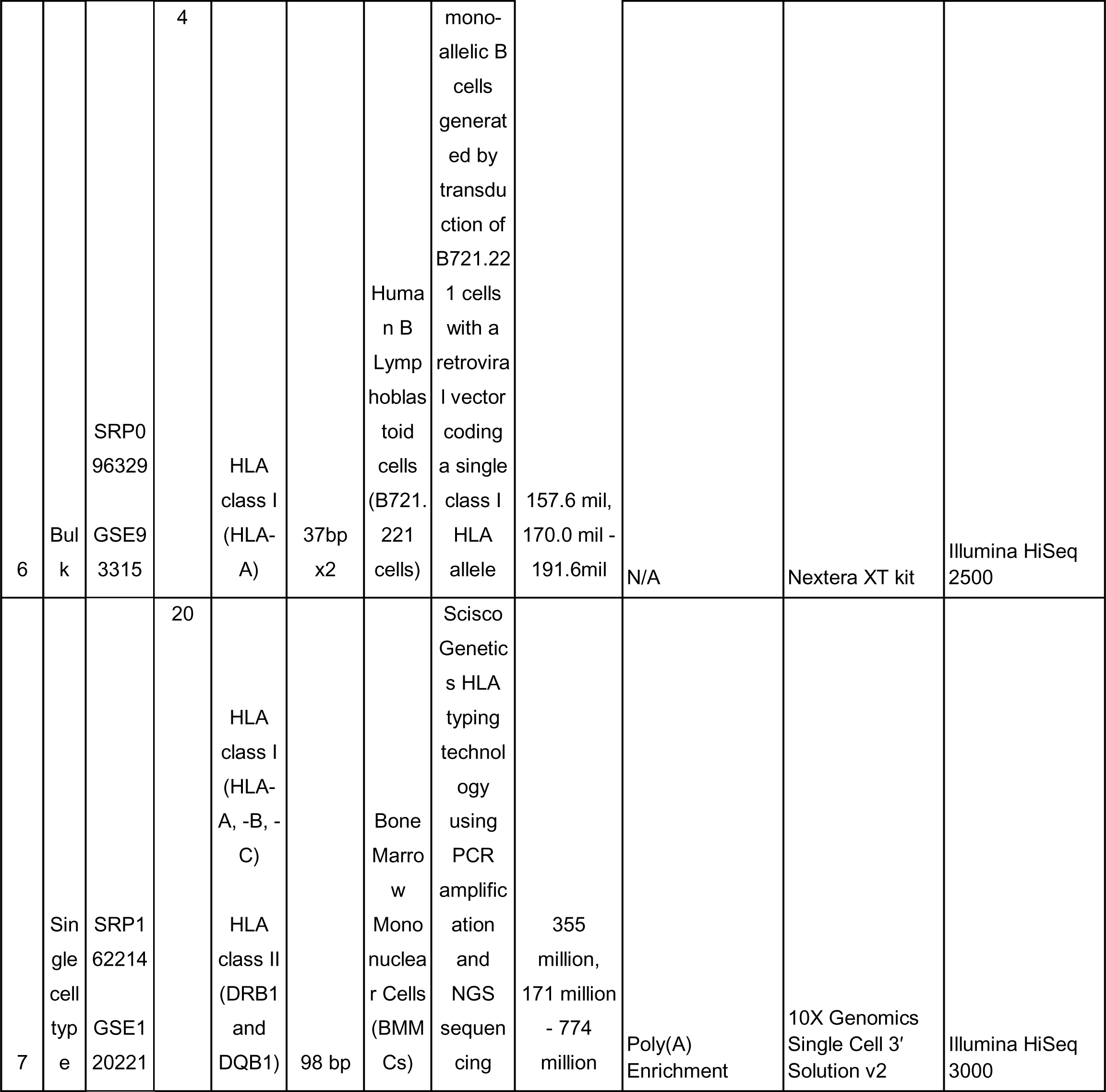
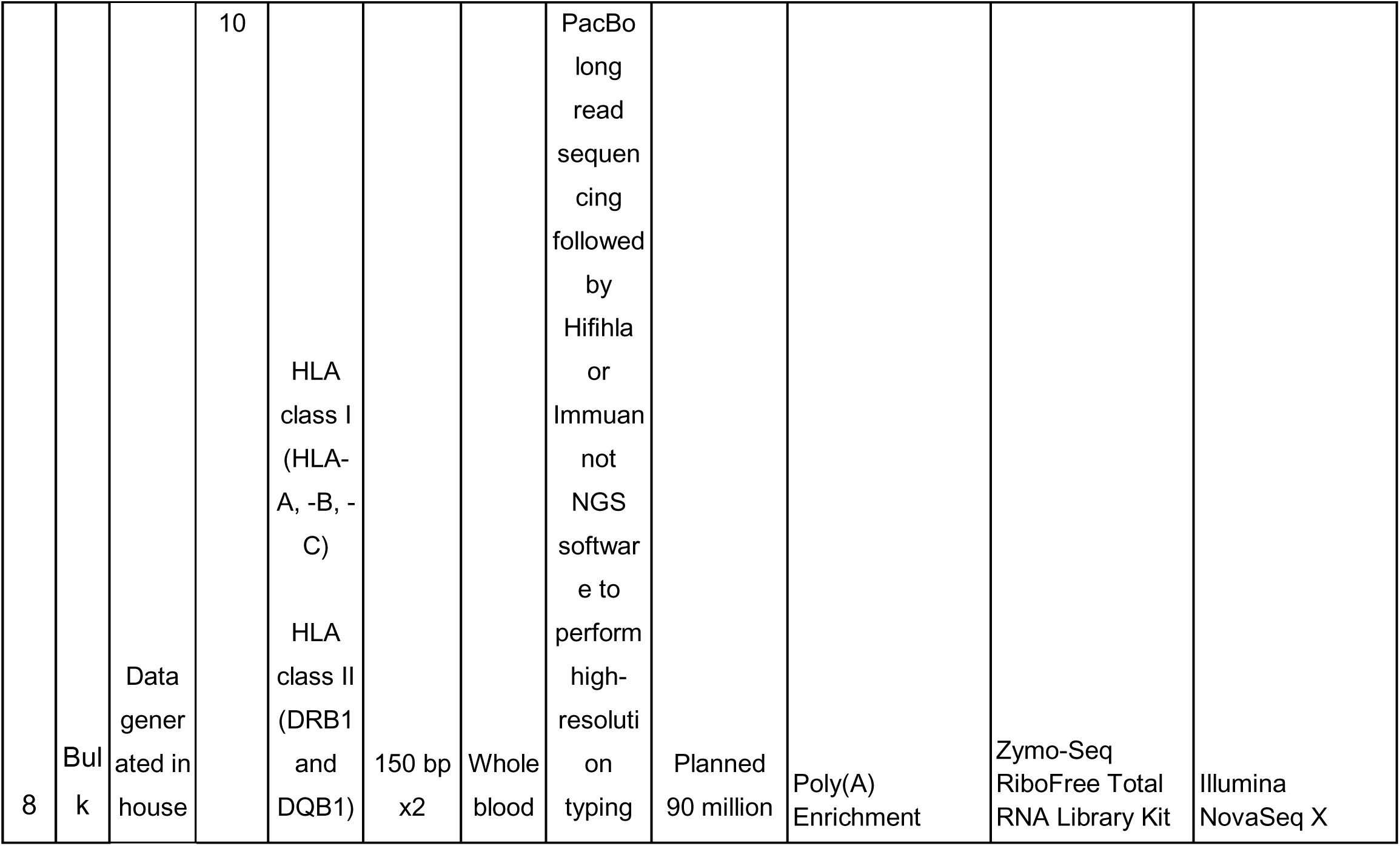
Overview of the gold standard datasets. “RNA-seq type” indicates whether the samples are bulk or single cell RNA-seq. “NCBI SRA Accession Number” indicates the unique identifier code for the complete dataset in the NCBI Gene Expression Omnibus record. “Number of samples” indicates the total number of samples in the dataset. “Genes with Gold standard HLA types” indicates the HLA loci for which there are gold standard HLA types determined for each dataset. “Read length” indicates the average number of base pairs per read, and “x2” indicates paired-end reads. “Tissue or cell type” indicates the type of cell or tissue that RNA was extracted from for RNA-seq. “Gold standard HLA typing method” indicates the method of determination of the specific allele for each gene of the HLA loci, to establish the HLA types gold standard. “Read count before extraction” indicates the number of reads in the raw samples prior to extraction of relevant reads for our benchmarking study (see “Preparation of RNA-seq input for HLA callers” subsection of Methods). “Library preparation method” and “Library preparation kit or custom protocol” briefly describe the library preparation and protocol used to perform RNA-seq for the samples of each dataset. “RNA-seq platform” describes the sequencing platform used. “N/A” is used to denote information not available in published literature.

All samples included annotated HLA genotypes at 5 loci, HLA-A, -B, -C, -DRB1, and -DQB1. The gold standard HLA types were determined via several protocols, including PCR-based sequence specific oligonucleotide probes (SSOP) (n=540), PCR-based sequence-specific primers (n=100), transduction of LCLs with a retroviral vector encoding for one or more HLA alleles (n=12), Scisco Technology novel genotyping technology (n=20), and PacBio novel genotyping technology (n=10).

Available alignment-based HLA callers that accept RNA-seq data were included in this benchmarking study, which include HLAminer (read alignment mode)^11^, seq2HLA^22^, HLAforest^23^, PHLAT^24^, OptiType^25^, HLA-VBseq^26^, HLA-HD^27^, HISAT-genotype^28^, arcasHLA^29^, HLApers^30^, RNA2HLA^31^, T1K^32^. Details on each tool can be found in Tables 2-3. We excluded HLAProfiler^33^ and bwakit^34^ from our analysis due to unresolvable errors during the installation or execution process (Table S3).

**Table 2:**
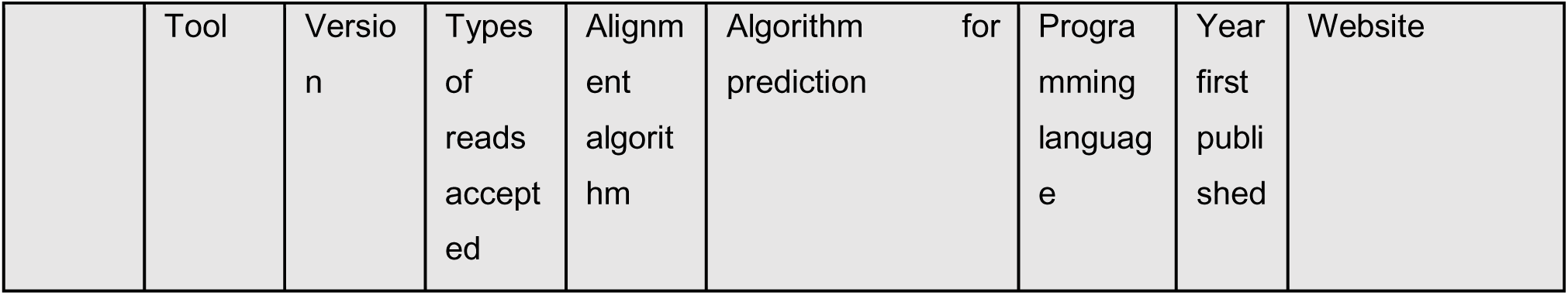
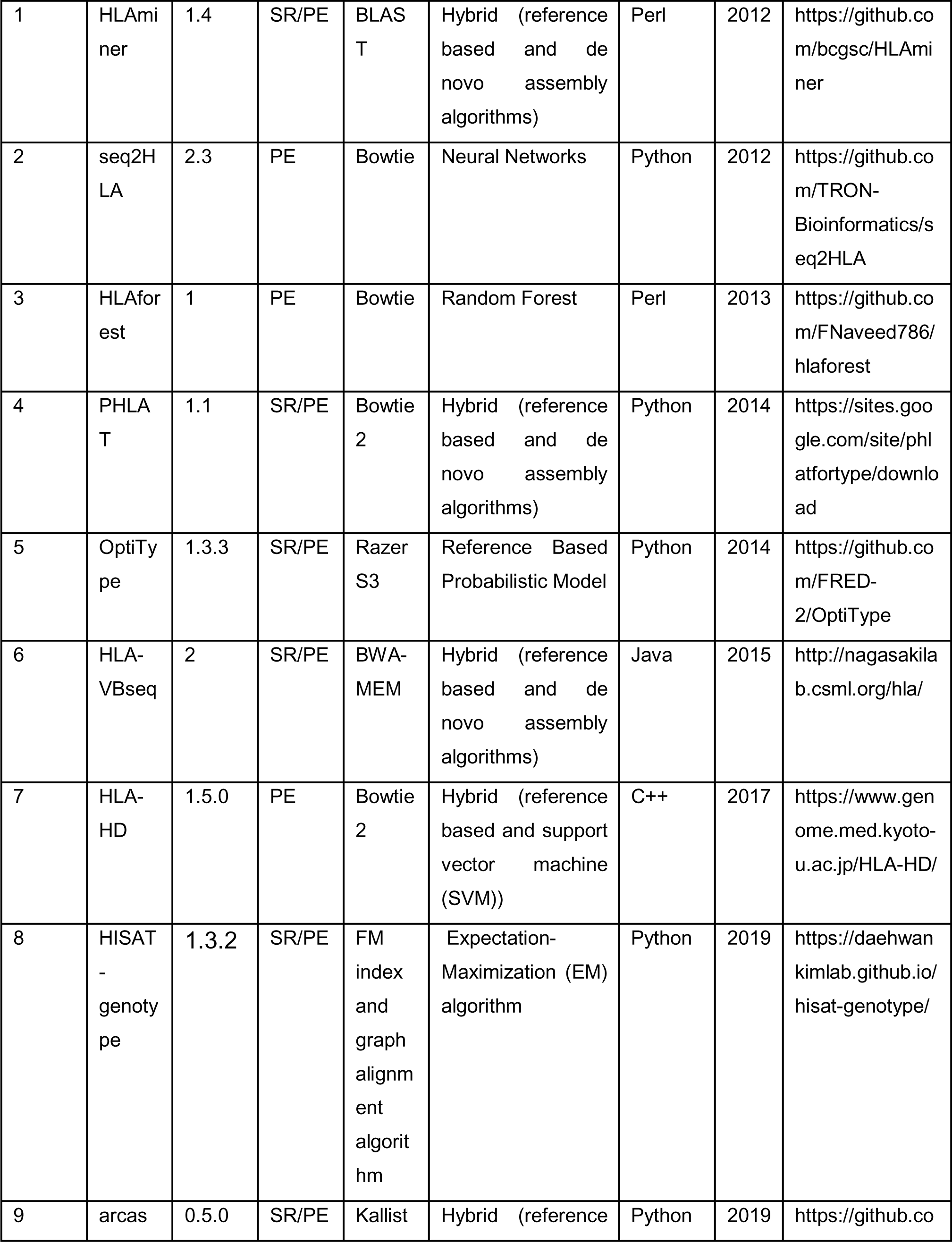
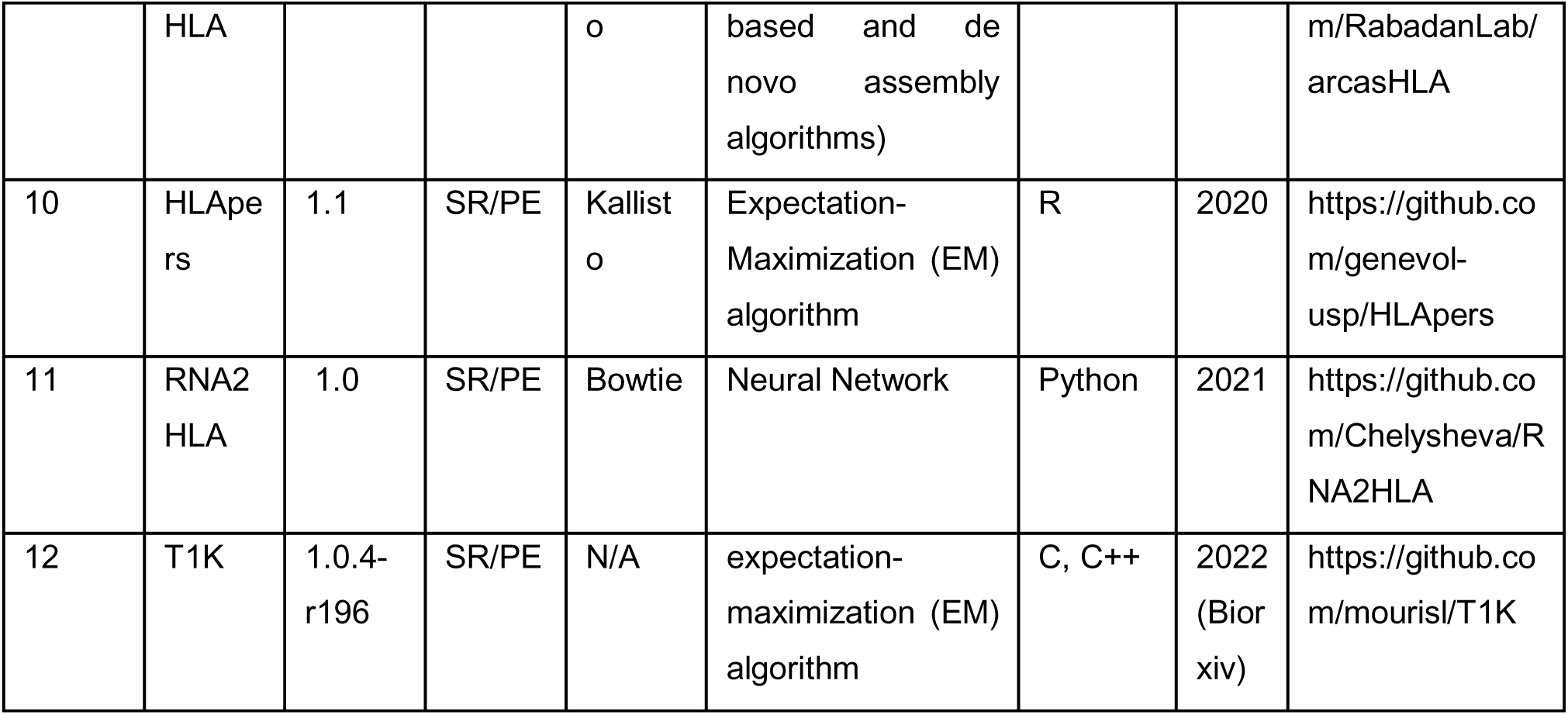
Overview of HLA callers included in this study. “Tool” indicates the names of each of the 12 HLA caller tools benchmarked. “Version” is the software version of the tool which was downloaded and installed for benchmarking. “Types of reads accepted”: SR indicates Single-Read, PE indicates Paired-End. “Alignment algorithm” indicates the algorithm used for alignment of reads to the reference genome. “Programming language” indicates the programming language in which most code for the HLA caller was written in. “Published year” is the year in which the tool and corresponding paper were published. “Website” indicates the GitHub repository link from which the software was downloaded. “N/A” is used to denote information not available in published literature.

**Table 3:**
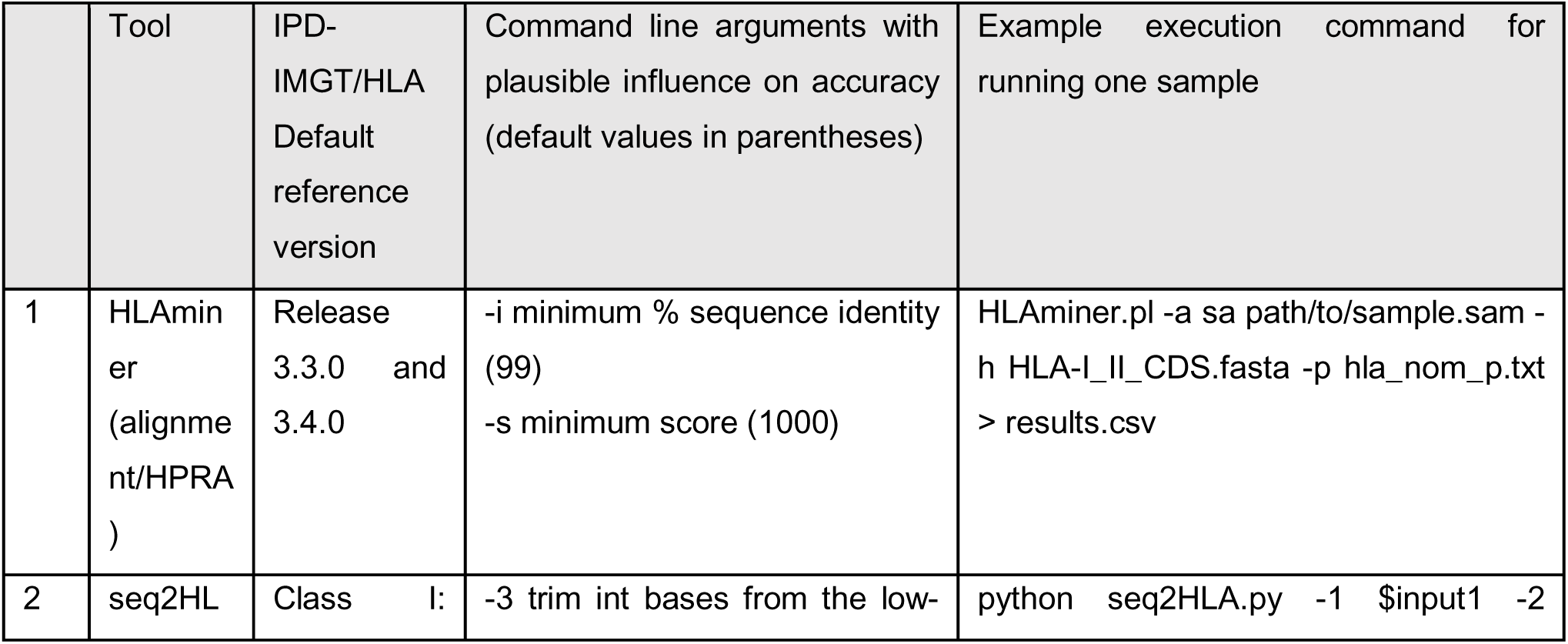
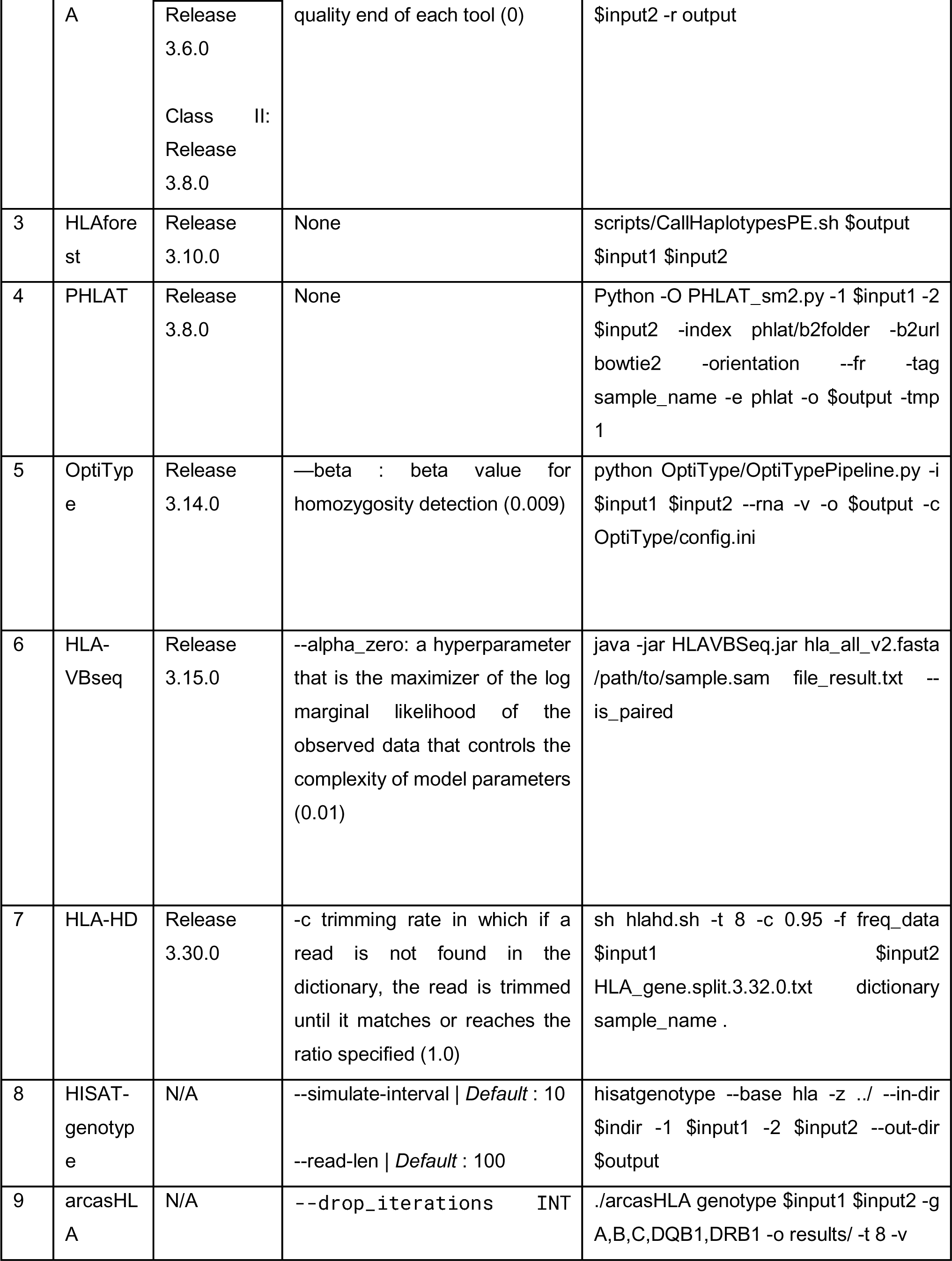
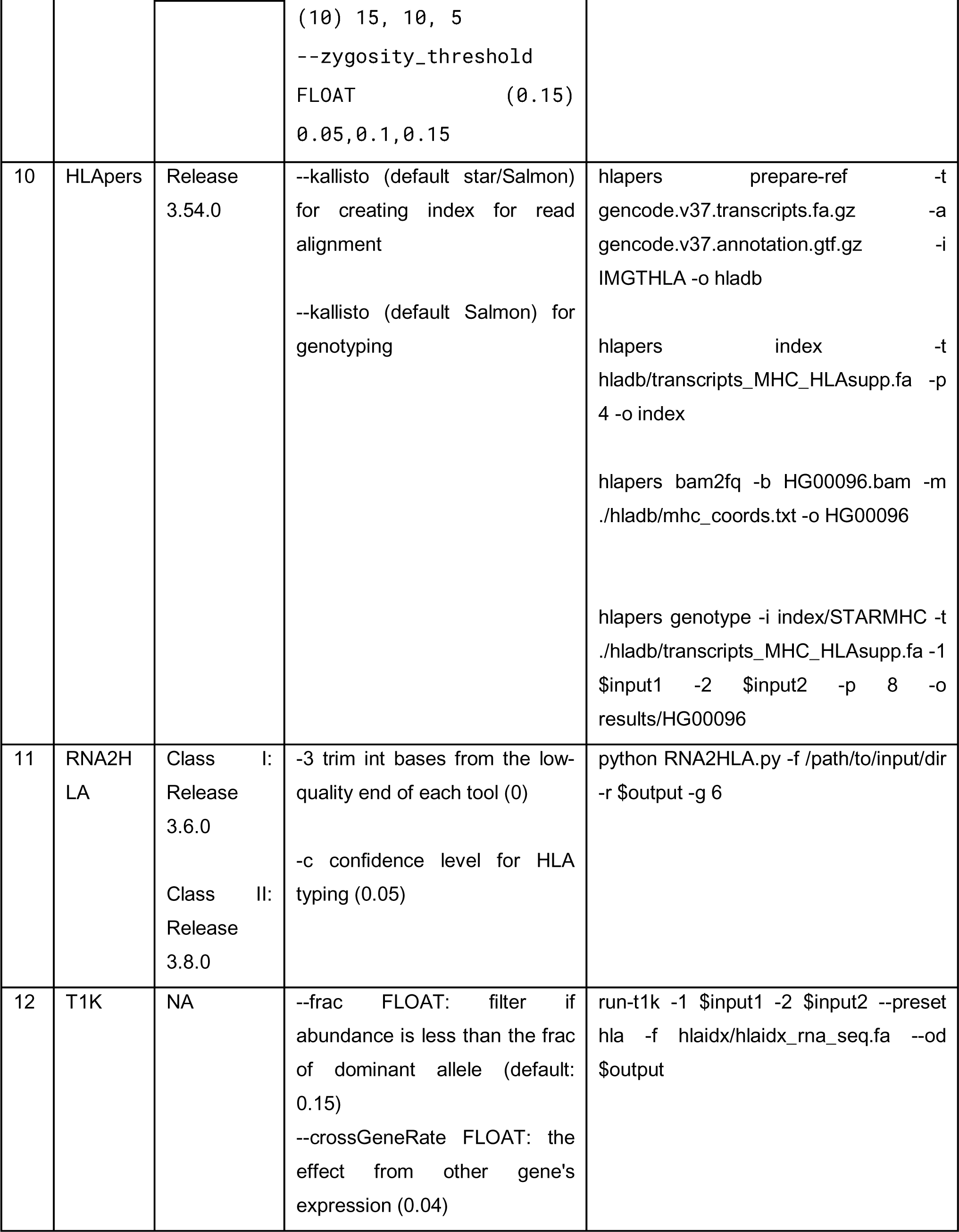
Commands used to install and execute the HLA typing tools to obtain HLA prediction results. “IPD-IMGT/HLA default Reference version” indicates the version of the software that is the default version. “Command line arguments with plausible influence on accuracy” include one or two parameters, when applicable, that influence the typing algorithm and that will be optimized as part of this study. We use “$input1” and “$input2” as placeholder names for the two paired-end fastq files, and “$output” as a placeholder name for the output directory, when applicable. “Example execution command for running one sample” lists the full command used to run one sample in the study. “N/A” is used to denote information not available in published literature.

### T1K has the greatest overall accuracy across Class I and Class II loci

At one-field resolution, the accuracy varies from 14.9% to 99.4% for Class I predictions, and 7.4% to 98.4% accuracy for Class II predictions. At two-field resolution, the accuracy is lower, with class I accuracy ranging from 6.4% to 98.3% accuracy, while Class II predictions ranging from 0% to 93.4% accuracy (Fig 3-4).

**Figure 3:**
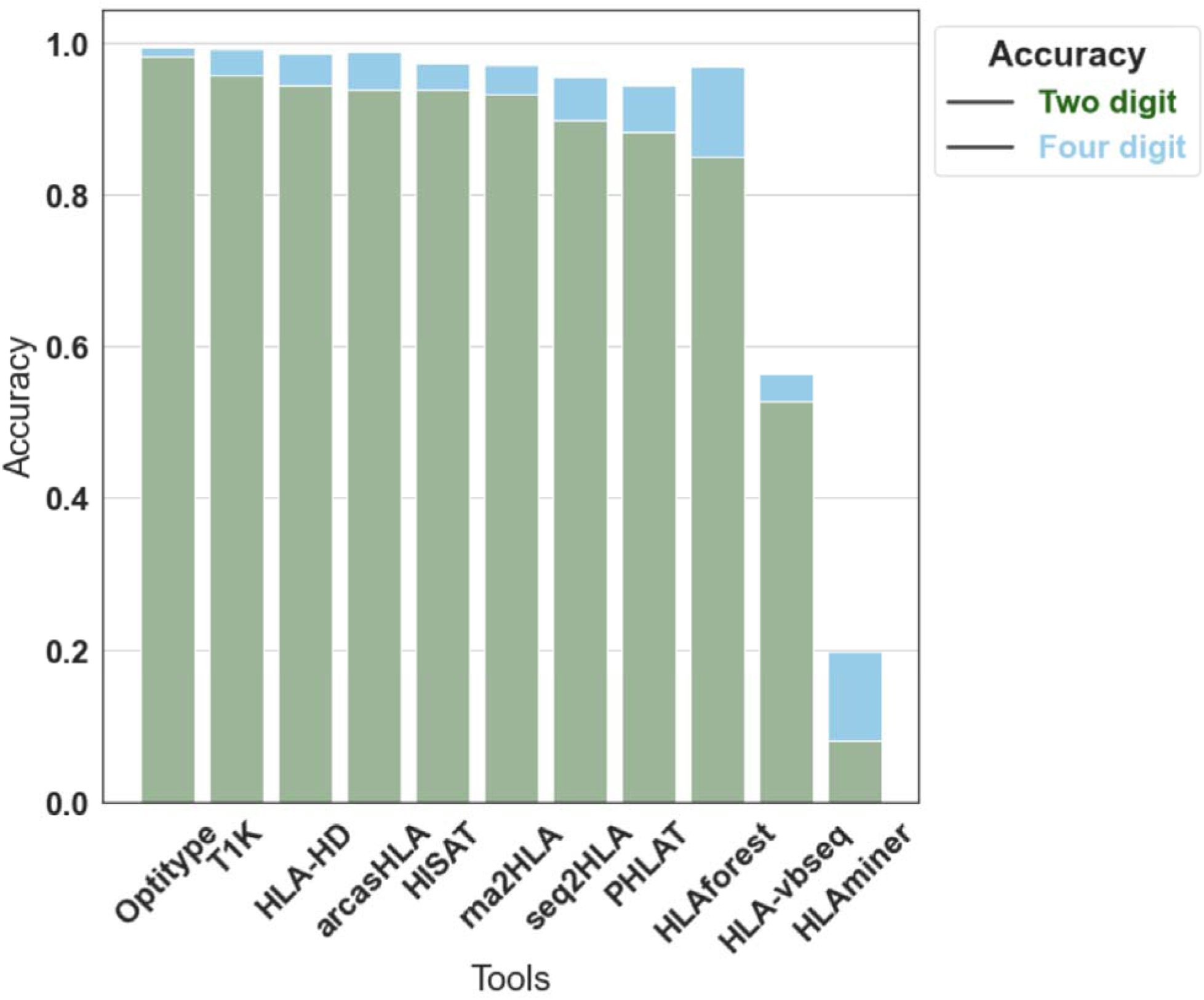
Accuracy of RNA-seq based HLA callers at one-field and two-field resolution. Tools are indicated on the y axis and accuracy is the y axis. Accuracy is measured on a scale of 0 to 1, where 1 indicates perfect accuracy while zero indicates that all imputations are incorrect. One-field resolution is represented in green and two-field resolution is represented in blue.

**Figure 4:**
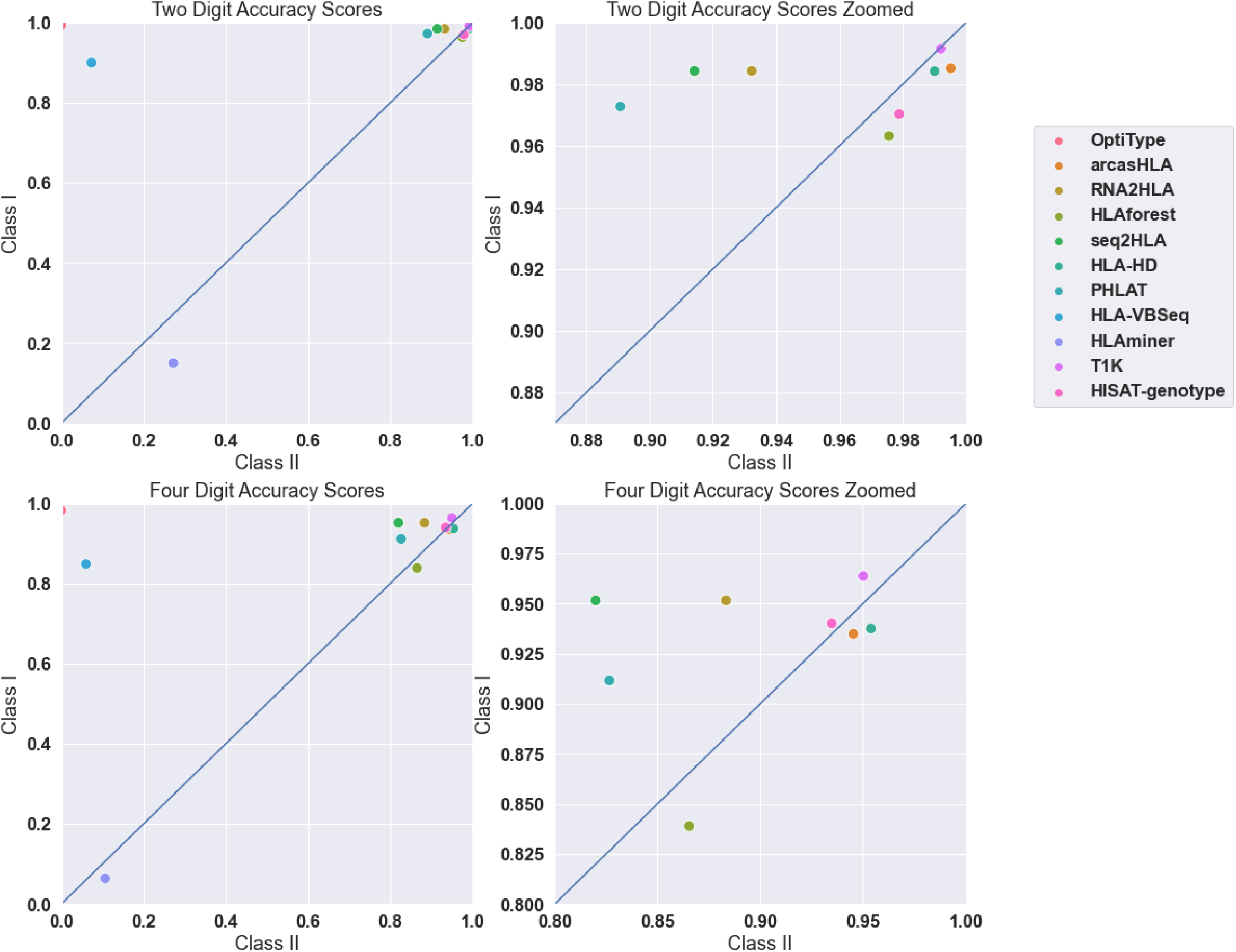
Accuracy of each tool to one and two-field accuracy, plotted with Class I accuracy against Class II accuracy.

At both one-field and two-field resolutions, for Class I alleles only, OptiType has the best performance in terms of accuracy for Class I alleles with scores of 99.4% and 98.3% respectively. For Class II alleles, T1K has the best accuracy performance with scores of 98.4% and 93.4%. Notably, T1K is able to make both Class I and Class II predictions with consistently high accuracy, whereas OptiType is only able to make class I predictions. HLA-VBseq and HLAMiner both struggle to make accurate predictions and have the worst performance for both Class I and Class II. These tools have a respective accuracy of 18.3% and 56.4% at one-field resolution, and 6.4% and 0% at two-field resolution. Exact accuracy values of each tool from 0 (fully inaccurate) to 1 (fully accurate) are reported in Table S2a-c.

### The HLA-B and HLA-DRB1 loci are most commonly mispredicted

We assessed each tool’s performance on each locus (A, B, C, DRB1, DQB1) to determine differences in accuracy across loci (Fig 5). We found that Class II underperforms compared to Class I, with the least accurate loci being HLA-DQB1 (70.1%) and HLA-DRB1 (75.5%) at one-field of resolution. Of the Class I loci, HLA-B performs the worst, with a 75.6% accuracy, whereas HLA-A and HLA-C both perform fairly well with accuracies of 89.3% and 88.8% respectively at two-field resolution.

**Figure 5:**
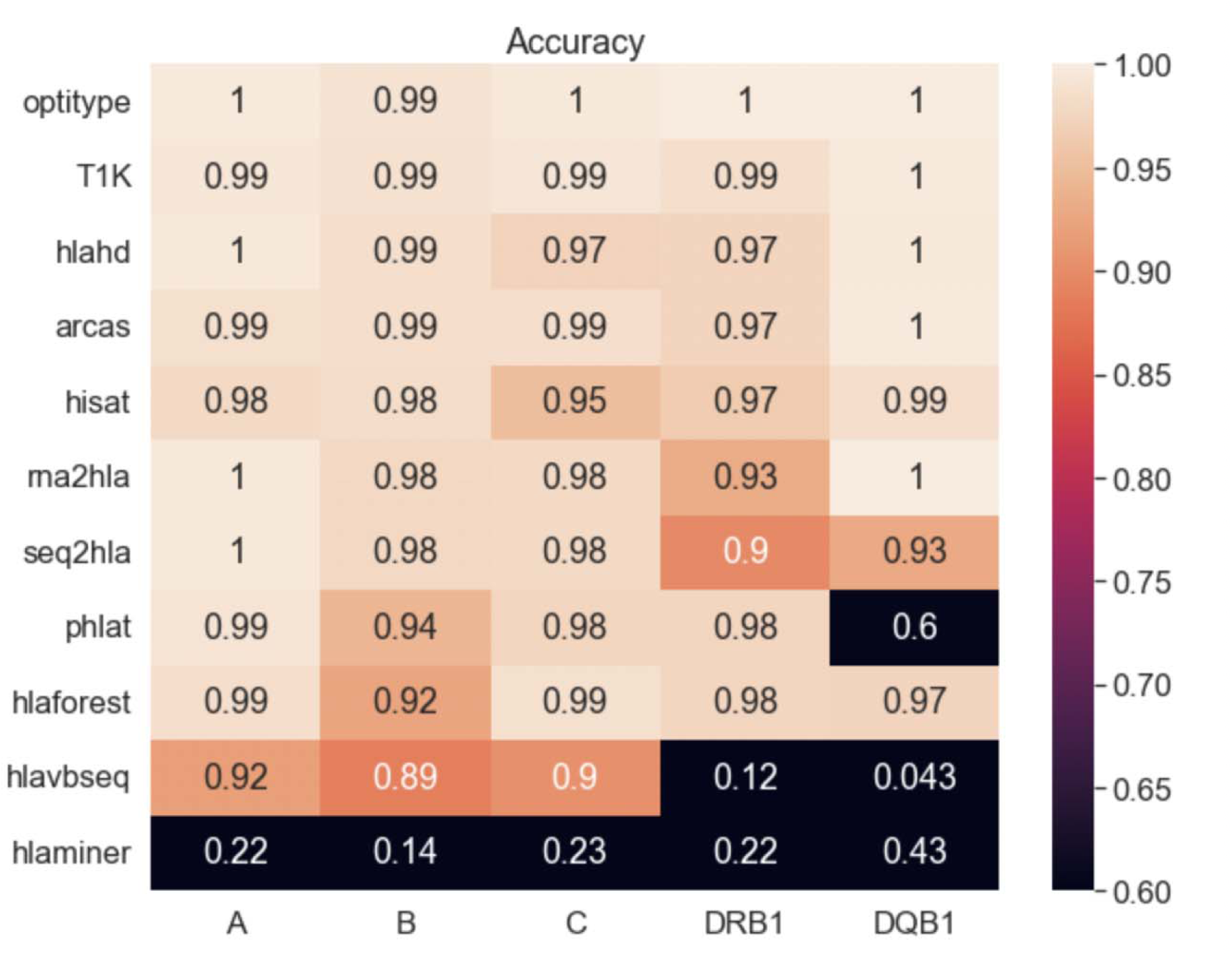
Number of Miscalled Alleles by Allele for Gold Standard Dataset 1. NOTE: OptiType can only predict Class I Alleles which is why it has no accuracy scores for Class II Alleles.

We additionally evaluated the percentages at which each individual allele in the gold standard datasets is mispredicted (Fig 6). Our results show that the Class II loci, particularly HLA-DQB1, is mispredicted at the highest ratio in comparison with alleles from other loci. Notably, we find that HLA-VBseq and HLAMiner are responsible for a majority of the Class II mispredictions, as removing these two tools from analysis decreased the significance of the difference in accuracies between Class I and Class II genes.

**Fig 6:**
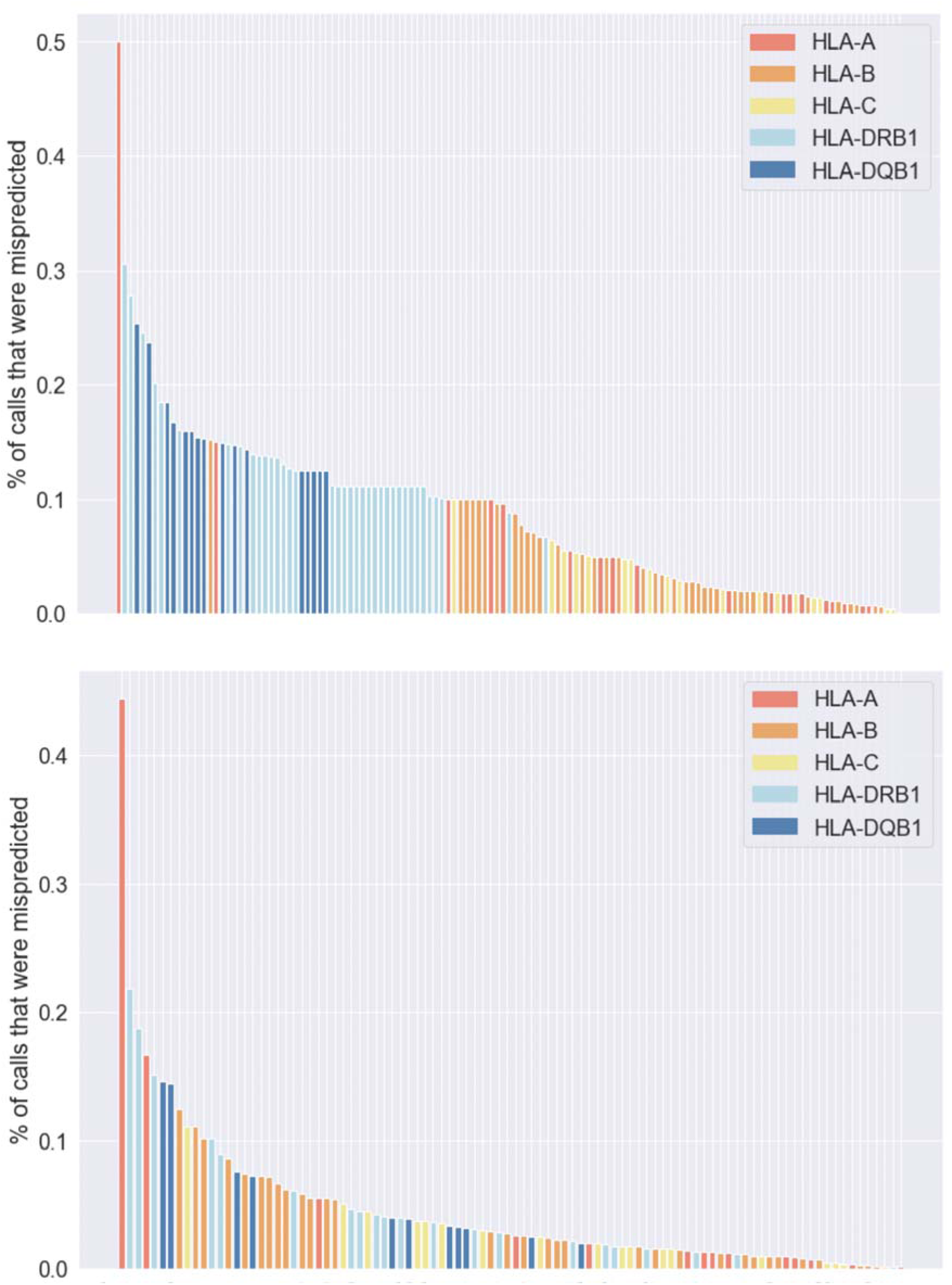
Misclassification rate of alleles. All alleles with height representing the % of mispredictions. Class I alleles are in orange whereas Class II alleles are in blue. Below is the same plot with 2 lowest accuracy tools (HLA-VBseq and HLAMiner) removed.

### Read lengths exhibit a moderate effect on prediction quality

From our preliminary analysis of 12 samples across Datasets 2 and 3, we find that increasing read length is associated with an increase in accuracy (Fig 7) for all tools except PHLAT, seq2HLA, and HLAMiner. A read length of 36 has an average accuracy of 63%, a read length of 51 has an average accuracy of 67% and a read length of 76 has an average accuracy of 77%.

**Fig 7:**
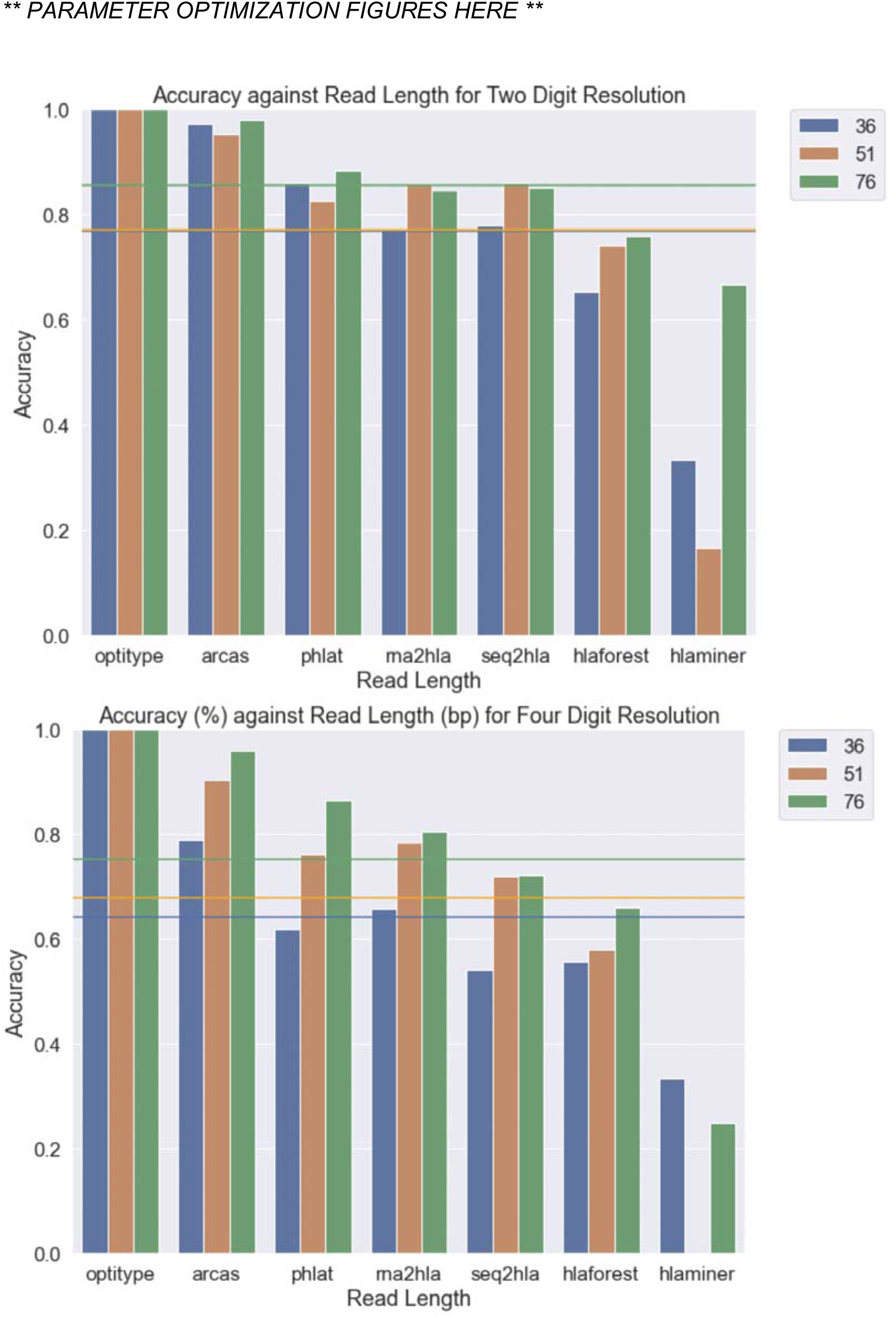
Accuracy of tools for three read lengths 36, 51, and 76. On the X-axis are the tools, and on the Y-axis is the numeric accuracy of the tool. Blue represents a read length of 36, orange represents a read length of 51, and green represents a read length of 76.

### All HLA callers have higher accuracy on European than African samples

We used all samples from Dataset 1 (n=490) to evaluate differences in performance between Europe and African ancestry groups. For each caller, we determined the accuracy at each HLA locus separately for European (n=423) and African (n=67) subsets. We observe the greatest disparity in accuracy in HLAforest^6^ (p=0.019) and PHLAT^7^ (p=0.017). Furthermore, we find that differences in accuracy are more pronounced at two-field resolution than they are at one-field resolution (Fig 8). At two-field of resolution, every single tool performed better on European than African ancestry groups. Although the disparity in some tools is not pronounced enough to be statistically significant, a binomial test demonstrates that the overall lower accuracy in African ancestry groups across all tools is significant (p=0.0039).

**Figure 8:**
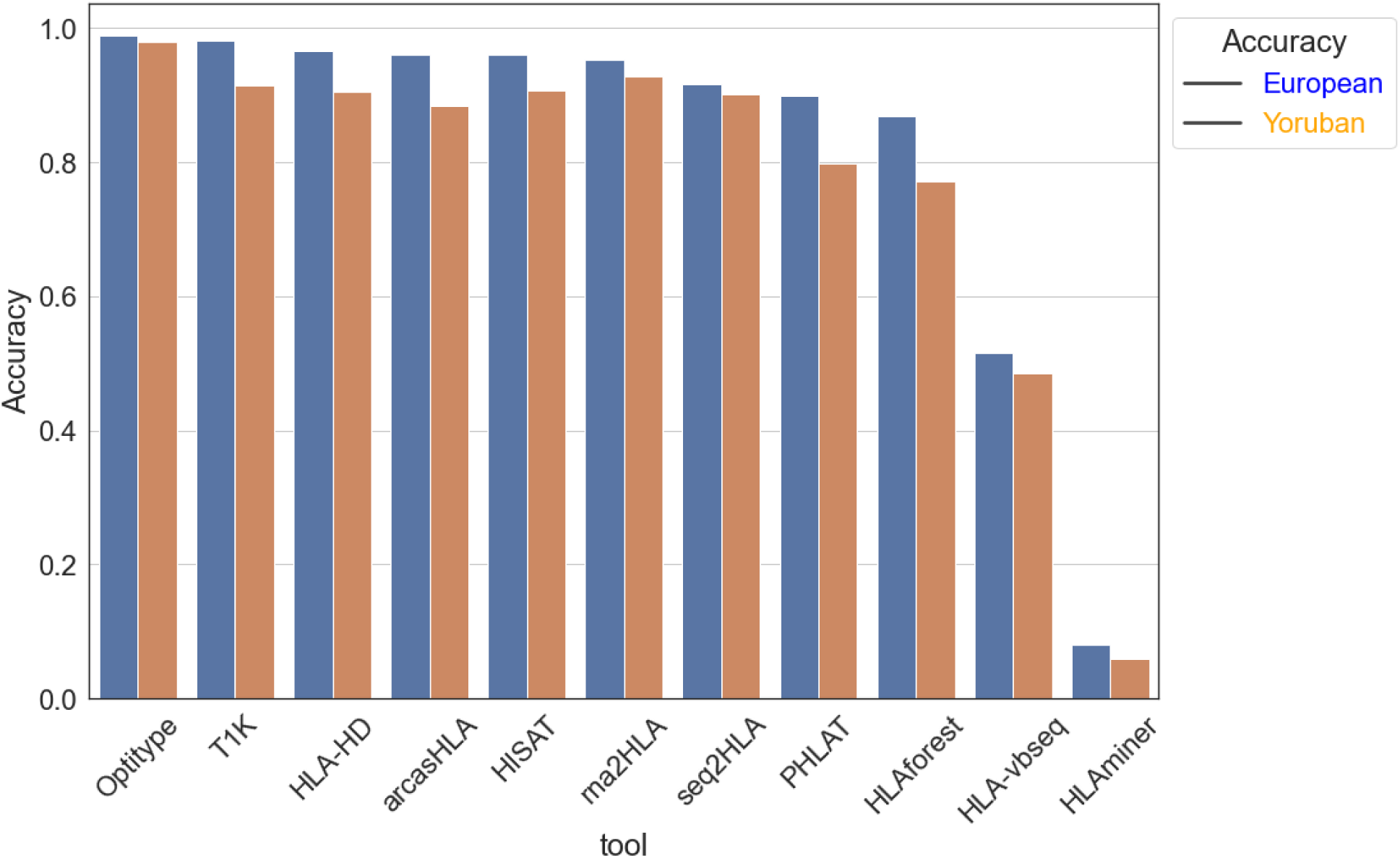
Accuracy between European and Yoruban samples for all tools. On the X-axis are the tools, and on the Y-axis are the numerical accuracies. Blue represents the European samples accuracy, whereas orange represents the Yoruban samples accuracy.

When we stratify the accuracy of each ancestry group by locus (Fig 9), we find that there is some variability in whether European or African subsets have greater accuracy for Class I alleles, but a vast majority of tools seem to be better at making accurate predictions for the European group for Class II alleles. The greatest discrepancy in accuracy out of the Class II genes is HLA-DQB1 in which the African subset accuracy is 2.7% lower than the European subset. Less extreme but still significant, between the Class I genes the greatest discrepancies in accuracy are observed in HLA-B where the average African subset accuracy is 1.3% lower than the European subset. Furthermore, for HLA-A the difference in average accuracy is 1.3% and for HLA-DRB1 it is 1.0%. HLA-C is the only loci where the African subset outperformed the European subset on accuracy, with a difference of 0.3 percentage points. We then isolated alleles and plotted the Europe against Africa misclassification rate for each allele, finding a slight skew towards higher misclassification rates for the Africa samples. We also notice a significantly greater misclassification rate for Class II alleles for both Europe and Africa alleles, which disappears with the removal of the tools HLA-VBseq and HLAMiner, again verifying that these two tools are insufficient for making accurate Class II imputations.

**Fig 9:**
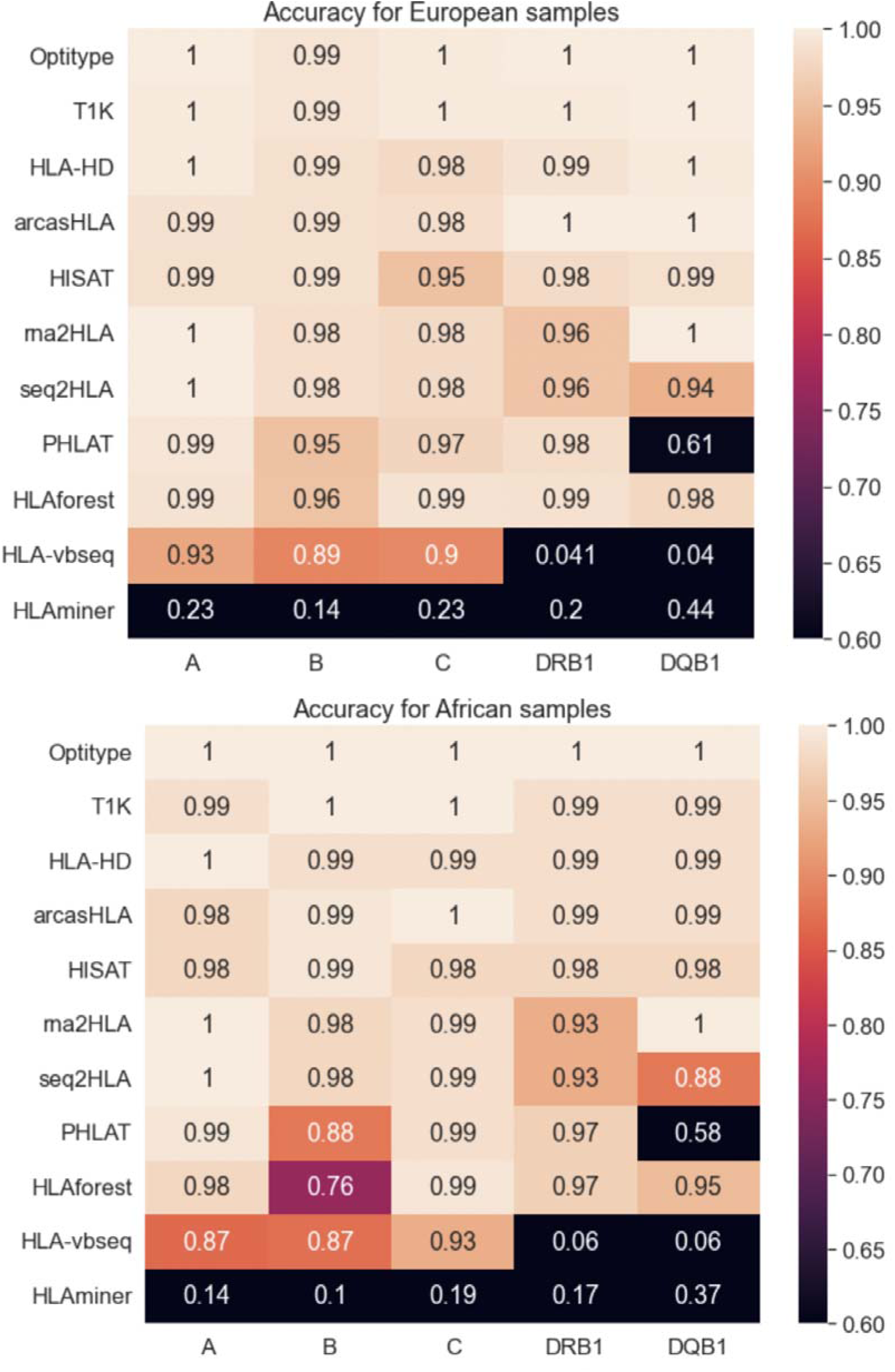
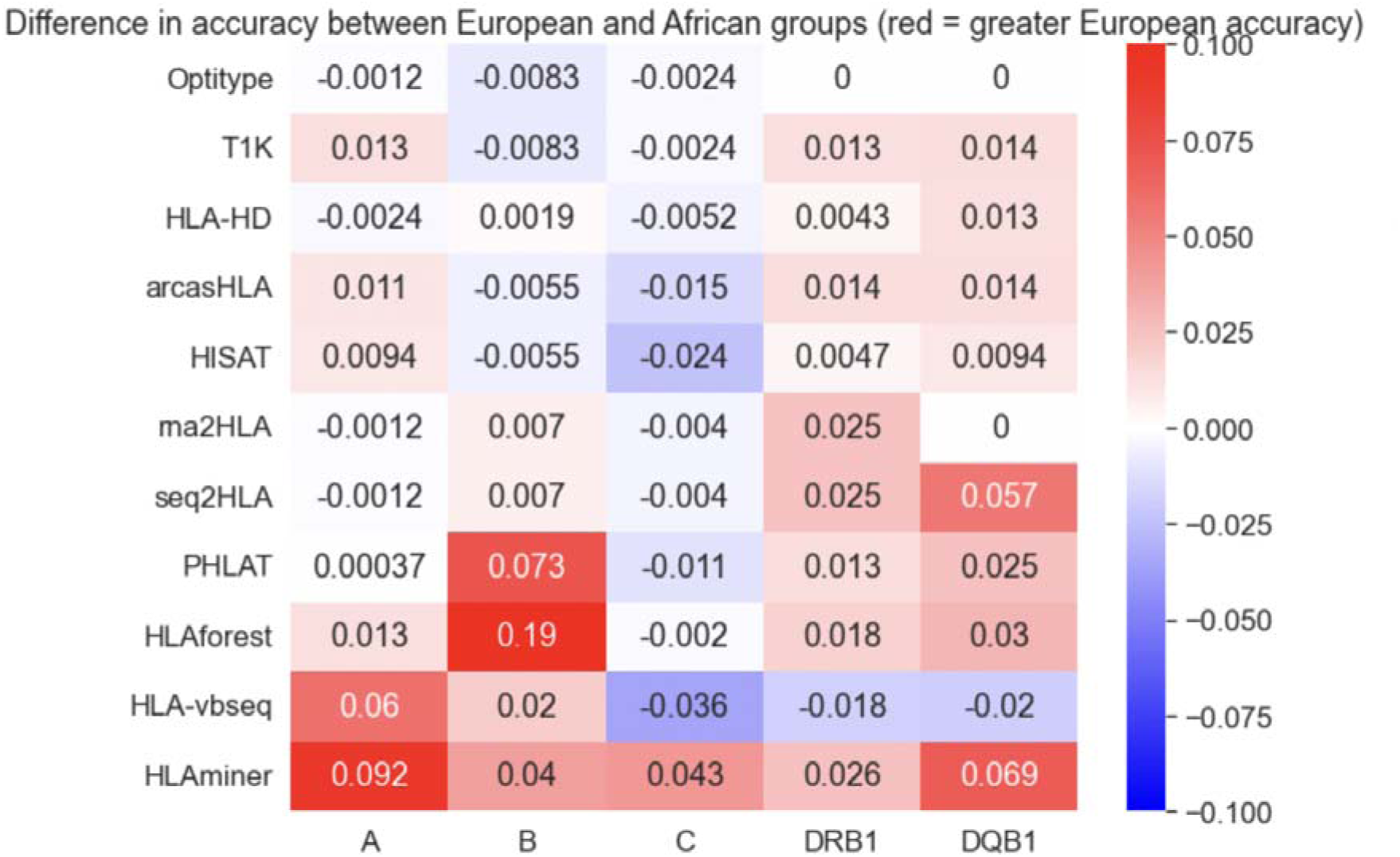
Accuracy of each tool for each of the 5 loci of interest HLA-A, -B, -C, -DRB1, and -DQB1. A) Accuracy of each tool at each locus for Yoruba samples (n=67). B) Accuracy of each tool at each locus for European samples (n=490). C) The difference in accuracy between the Yoruban and European subsets. Blue represents tools and loci for which Yoruban samples have higher accuracy, while red represents tools and loci for which European samples have higher accuracy.

**Figure 10:**
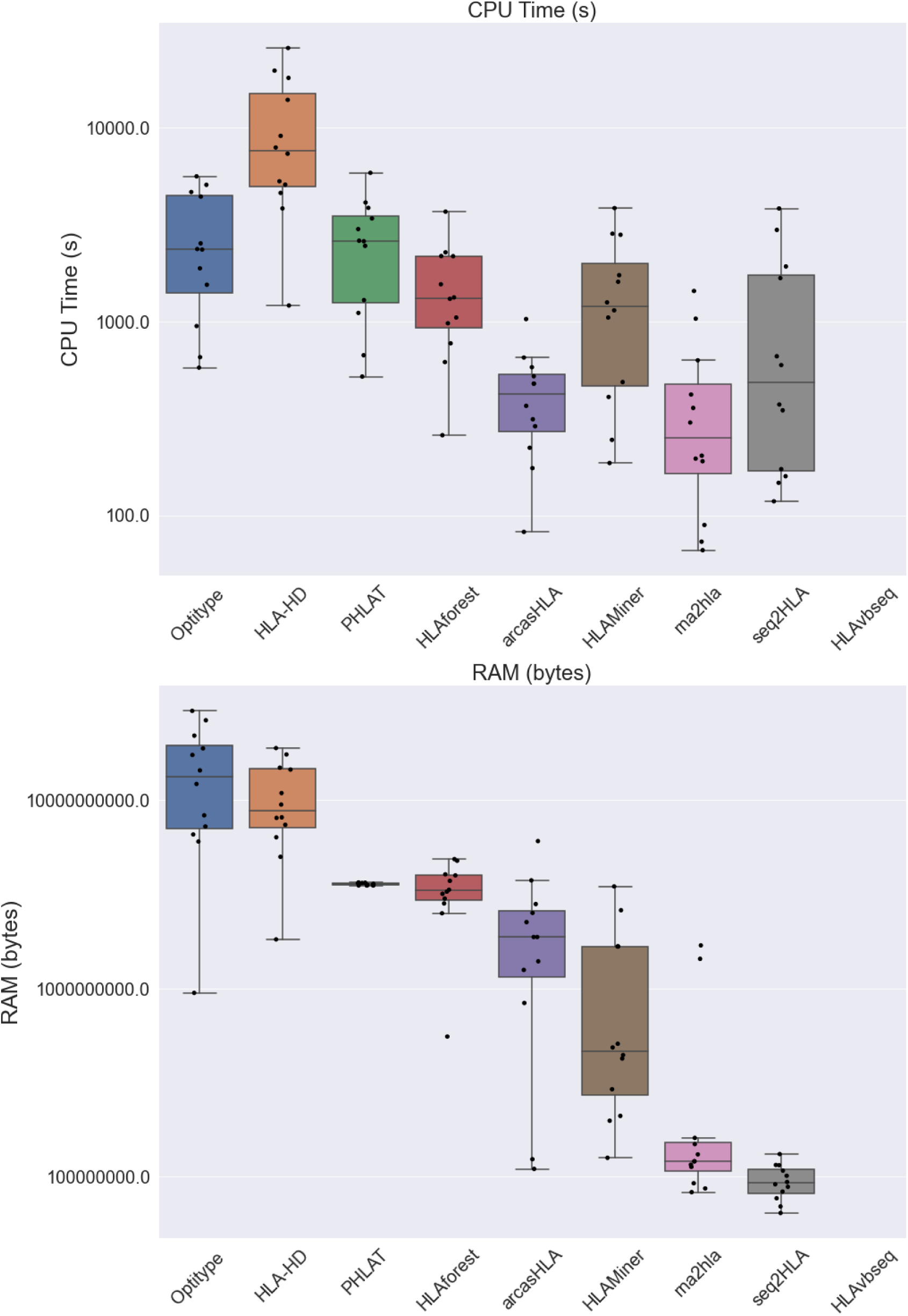
Memory and CPU, by caller, for calling of alleles for a subset of samples in Datasets 1-6 (n=12).

### Computational resource needs vary greatly across the HLA callers

We measured the CPU time and RAM usage of each caller for two samples per dataset to assess computational expensiveness (Fig 3a-b). We find great variation in the computational resources necessary to run the tools, with RAM ranging from the order of 100 kB to 100 GB, and CPU time ranging from approximately 10 minutes to 3 hours. While there is often a trade off between accuracy and computational resources required by each tool, we observe that RNA2HLA yielded the best balance between high accuracy (93.3%) at low computational expense (average 0.36 GB RAM amd 6.9 min CPU time), making it suitable for HLA typing with large scale studies. In contrast, the best accuracy tool, OptiType, requires an average of 14.2 GB RAM and 45.3 min CPU time per sample.

## Discussion

Our rigorous assessment of HLA callers highlights the advantages and limitations of computational HLA typing algorithms across several Class I and Class II loci and various sequencing parameters. Altogether, this study offers crucial, up-to-date information for researchers regarding appropriate choice of tool for an HLA typing from RNA-seq data. Furthermore, our study demonstrates the potential for RNA-seq based HLA typing tools to outperform traditional methods as a scalable and cost-effective method for imputing HLA typing. Crucially, our comparison of the performance of HLA callers across different ancestry groups highlights the pressing need to develop improved HLA typing algorithms and databases that can accurately account for the distinct genetic variations present in non-European populations.

In comparison to the previous RNA-seq benchmarking study^21^, we found highly consistent results with the same tools performing well in their study, OptiType and arcasHLA performing similarly well and HLA-VBseq struggling to make accurate imputations. We also found that more recently developed tools perform similarly well to prior tools, without many significant improvements to accuracy in the past few years. For instance, Optitype remains the most accurate tool for Class I genotypes. All tools performed much better on Class I than on Class II genes, with Class II accuracy being much lower compared to the current gold standard HLA typing methods.

Our study is the first to evaluate the ability for HLA callers to impute HLA types from specifically RNA-seq data. We find that all tools, except HLA-VBseq and HLAMiner, are capable of making Class I predictions to two-field (clinical) resolution to 82.6% - 98.3% accuracy when compared with the gold standard method. We find that there are few tools that are able to make consistently accurate predictions across all loci and classes, and we believe that arcasHLA is the overall best tool as it performs with a consistently high 93.4% accuracy on Class I and Class II. In addition to arcasHLA, we also recommend RNA2HLA as a tool for high quality predictions at relatively low computational expense. Our study is also the first to explore the effect of read length and ancestry on the imputation quality of the callers. We found that read length does not have a substantial effect on accuracy. The tools are more accurate at making predictions for individuals of European ancestry than those of African ancestry.

In the future, we plan to complete the read length analysis and data collection for the CPU and RAM usage of the tools as we are currently missing some data. We also plan to determine the effect of coverage on the accuracy of the HLA typing algorithms. We also plan to determine the concordance in predictions across various samples, to evaluate whether the algorithms have the tendency to predict certain alleles. Once identifying patterns of misclassifications across tools, we also hope to analyze the sequence similarity of the allele with the misclassification to establish a hypothesis for why these errors occur. One final step we hope to complete is running tools with various custom parameters and arguments to determine the best set of parameters for each tool, rather than simply the default parameters like we are currently doing.

We envision that future HLA callers will accurately impute HLA genotypes for both Class I and Class II genes to two-field resolution, thereby offering clinicians and researchers a well performing and scalable option for HLA typing with WGS data. Current methods show an inability to make accurate Class II predictions, and discrepancies in prediction quality across different ancestral groups, and addressing these challenges will make HLA callers a viable choice for HLA typing studies. One potential limitation is that our CPU/RAM was collected from jobs submitted to a remote high performance computing cluster, and hence may not be completely controlled because it might be dependent on resource availability of the cluster at the time we ran each sample. We mitigated this limitation by running each sample 10 times and taking the average to be used for our analysis. We hope this study offers crucial information for researchers regarding appropriate choices of methods for conducting HLA typing with RNA-seq data.

## Methods

### Bulk RNA-Seq data

We will retrieve data from 6 publicly available bulk RNA-seq datasets (ERP001942^35^, SRP298704^36^, ERP000101^37^, SRP099568^38^, SRP198497^39^, SRP096329^40^) to assess the performance of the tools (Table 1). From the datasets we will select all samples with annotated HLA genotypes determined via gold standard methods, amounting to a total of 652 samples collected from lymphoblastoid cell lines (LCLs) (n=552) and peripheral blood mononuclear cells (PMBCs) (n=100). Altogether, these datasets represent a diverse set of samples spanning 5 1000-genomes project populations (Northern Europeans in Utah, Finnish, British in England and Scotland, Tuscan in Italy, Yoruban), 4 sequencing platforms (Illumina HiSeq 2000, Illumina HiSeq 2500, Illumina Genome Analyzer II, Illumina NextSeq 550), read counts ranging from 24.5 million to 337.8 million reads, and read lengths ranging from 36bp to 126bp.

We will also generate RNA-seq reads for samples corresponding to the D8 benchmarking dataset, consisting of whole blood samples from healthy donors sequenced with RNA-seq (Zymo Research). cDNA libraries were constructed via Poly-A selection, and libraries were sequenced on the Illumina NovaSeq X platform at 90 million reads and 2×150bp read length.

### scRNA-seq data

Additionally, we will download a 3’-based single cell RNA-seq dataset by the accession number SRP162214^41^ (Table 1) containing 20 samples collected from bone marrow mononuclear cells (BMMCs) from healthy donors. Droplet-based scRNA-Seq was performed using 10X Genomics Single Cell 3′ Solution v2 and libraries were sequenced on the Illumina HiSeq 3000 platform. The samples have a mean read count of 355 million reads, and reads are 98 bp in length.

### Preparation of RNA-seq input for HLA callers

We will download, align, and extract the RNA-seq reads to samples serving as inputs to the HLA callers. First, we use the SRA-Toolkit v2.11.0 prefetch and fasterq-dump commands with default parameters to download the raw RNA-seq reads as FASTQ files from the NCBI SRA archive. Then, we will use the fq v0.11.0 tool, fq lint command, to check the integrity and validity of the fastq files.

We will then use STAR aligner v2.7.0e to align the high quality reads of each sample to the GRCh38 reference using the STAR command with default options, except ‘--genomeDir’ to specify the path to the GRCh38 reference, ‘--readFilesIn’ to specify the paths to the two paired end fastq files, ‘--genomeDir’ argument to specify the location of the file containing the genome indices, “--outReadsUnmapped Fastx” to retain all unmapped reads, and ‘--outSAMtype BAM SortedByCoordinate’ indicates the output files to be sorted BAM files.

From the aligned BAM files, we will extract the chromosome 6 regions using Samtools v1.19, with default parameters unless otherwise specified. We will first use the Samtools index function to produce indexed BAI files from the BAM-format samples for fast random access. Then, will invoke the samtools view command with the ‘chr6’ parameter to extract the reads aligning to chromosome 6. Then, we will invoke the samtoolsview -q 30 to isolate only high quality reads with >30 mean sequence quality. Finally, we will invoke samtools view with the -b argument to convert the extracted and filtered samples to bam files, and the bam2fq command to convert those bam files back to FASTQ files. These prepared FASTQ samples will serve as the final inputs for all HLA callers.

### HLA definition and nomenclature

To specify each HLA allele type, will use the HLA nomenclature system defined by the World Health Organization Committee for Factors of the HLA System in 2010. This system uses a four-part naming system: the HLA Prefix, the gene, the field groups, and the suffix letter as shown in Figure 2. The first part, or the HLA Prefix, is denoted by the string “HLA-” and is universal across all alleles in the region. The second part denotes the name of the locus. The third part specifies the field groups, which are divided into 4 fields of increasing resolution with each field separated by a colon. Field 1 represents the allele group, or specifically what serological antigen type is present for this particular allele. Field 2, which is typically 2 or 3 digits long, represents the specific HLA protein. Field 3 represents allele variants with synonymous or intronic differences. Field 4 designates whether the allele is a single or multiple nucleotide polymorphism in the non-coding region of the gene. Finally, the suffix letter denotes differences in expression of the allele as described in Table S1.

One-field resolution refers to the allele name considering only up to the first field of the third part (i.e., HLA-A*02). Two-field resolution is the clinically relevant resolution and refers to the allele name considering only up to the second field of the third part (i.e., HLA-A*02:10). This nomenclature system categorizes each HLA type in increasing specificity, with the most general classifications to the left and most specific classifications to the right. As such, one-field resolution may also be referred to as low resolution, and two-field resolution may be referred to as high resolution or clinically relevant resolution. For the purposes of this benchmarking study, we will be assessing performance at the one-field and two-field resolutions.

### Gold Standard HLA genotyping via PCR-based methods

Annotated gold standard HLA types of samples in Datasets 1-6 were determined via laboratory PCR-based methods summarized in Table 1. For dataset 1, HLA types were found via PCR-based sequence-specific oligonucleotide probes (SSOP method) which uses PCR to amplify a specific region of the HLA gene, and then uses sequence-specific oligonucleotide probes that will detect and differentiate specific alleles or variants in that region based on their complementary binding. Dataset 2 and 4’s HLA types were found using PCR-based sequence-specific primers (SSP method) which amplifies the specific regions of DNA using primers that are designed specifically to bind to and amplify particular alleles of HLA genes. Dataset 3’s HLA types were determined using PCR-based sequence-specific oligonucleotide probes (SSOP method). Dataset 5 and 6 were created from cells from the HLA Class I-deficient B721.221 cell line transduced with one or two Class I HLA allele-expressing cDNA vectors.

It should be noted that gold standard datasets D1-D6 contain HLA types determined via laboratory-based HLA typing methods (PCR-SSP and PCR-SSOP) which are unable to fully resolve HLA types, resulting in phase ambiguities. Dataset 1, representing 490 (approximately 70%) of our samples, do contain original data with all phase ambiguities reported. But for Datasets 2-6, the available data contains samples with no reported allele ambiguities. Unfortunately, those studies did not publish details on the gold standard HLA typing to determine whether these samples simply had no loci with ambiguities, or whether all ambiguities were resolved prior to publication of the data.

We have formatted the gold standard HLA types for each dataset into a standard format csv where rows represent the samples and columns represent the loci(s) of interest (HLA-A, HLA-B, HLA-C, HLA-DRB1, HLA-DQB1) for which gold standard allele data are available. The csv files have been made available at https://github.com/Mangul-Lab-USC/HLA_benchmark.

### High-resolution HLA genotyping using Scisco HLA genotyping to produce a highly accurate gold standard

Gold standard HLA types in Dataset 7 contain samples typed via high-resolution Scisco HLA genotyping technology to produce highly accurate gold standard across 20 samples. Scisco HLA genotyping utilizes a 2-stage protocol involving amplicon-based PCR amplification of genomic DNA, followed by sequencing on the Illumina MiSeq platform and analysis using the ScisCloud software. PCR allows for complete coverage of all exons and sufficient intron sequences in order to detect all known intron encoded null alleles, Samples typed using this technology have unambiguous allele assignments to three fields of resolution. We have formatted the gold standard HLA types for Dataset 7 into a standard format csv where rows represent the samples and columns represent the loci(s) of interest (HLA-A, HLA-B, HLA-C, HLA-DRB1, HLA-DQB1) for which gold standard allele data are available. The csv files have been made available at https://github.com/Mangul-Lab-USC/HLA_benchmark.

### High-resolution HLA genotyping using long-read technologies to produce a highly accurate gold standard

We will generate a new dataset, labeled Dataset 8, with gold standard for 10 samples using high-resolution HLA genotyping from PacBio, which will produce a highly accurate gold standard for the 10 samples. We will perform the high-resolution gold standard HLA genotypes by performing whole genome sequencing at 30x coverage will be performed on the PacBio Revio long-read sequencing machine, followed by the Immuanot (https://github.com/YingZhou001/Immuannot) tool recommended by PacBio sequencing specialists, to generate four-field, full-resolution, fully-phased HLA calls. We will format the gold standard HLA types for Dataset 8 into a standard format csv where rows represent the samples and columns represent the loci(s) of interest (HLA-A, HLA-B, HLA-C, HLA-DRB1, HLA-DQB1) for which gold standard allele data are available, which will be made available at https://github.com/Mangul-Lab-USC/HLA_benchmark.

### HLA caller choice and installation

We plan to benchmark all existing alignment-based HLA callers capable of imputing HLA types from RNA-seq data. With these prerequisites, we identified 12 tools to be included in this study, namely HLAminer^11^, seq2HLA^22^, HLAforest^23^, PHLAT^24^, OptiType^25^, HLA-VBseq^26^, HLA-HD^27^, HISAT-genotype^28^, arcasHLA^28^, HLApers^30^, RNA2HLA^31^, T1K^32^ (Table 1). We will download each tool from their respective repository onto the USC CARC Discovery Cluster, closely following all available documentation and recommendations for each tool. If testing suites are provided, we will also run the tools on the testing suites and manually verify that all outputs are as expected. Furthermore, we will email the developers of each tool to verify our process of installation and for recommendations on best practices for installation and running. If any issues arise during the installation process, we will open an issue in the GitHub repository for the tool. After corresponding with developers and opening an issue on GitHub, if we are unable to successively resolve the installation errors then we will report the error in our results and exclude the tool from the study.

### Run HLA profiling methods

After installation, we will run each HLA caller by submitting bash scripted jobs to the SLURM job scheduler on the USC CARC Discovery cluster. We will run all tools on the first 6 datasets, and all tools able to accept single-read RNA-seq samples on the Dataset 7. We will run each tool on two versions of inputs, including the raw RNA-seq files, files containing only reads mapping to Chr6, and files containing only reads mapping to the Classical HLA region. Additionally, we will run tools on 20 samples from Dataset 1 that contain reads mapped to the Classical HLA region as well as all unmapped reads, to investigate the effect of considering unmapped reads on the accuracy of each HLA typing algorithm. Furthermore, we will run each tool on default parameters as well as conducting a sweep of all specifiable command line arguments to determine the optimum parameters for each tool (see “Parameter Optimization” section of Methods). Information on parameters and commands to run each tool can be found in Table 3. We will consult with the developers of each tool during the installation and running process to ensure that each tool is operated as designed.

### Parameter Optimization

All tools except HLAforest and PHLAT are configured with customizable command line arguments with a plausible influence on accuracy. For each such tool, we will explore the effect of alternative parameter values on accuracy and determine the optimal set of parameters, with a summary given in Table 3. To specify, for HLAminer we plan to test the effect of adjusting -i between values of 90 and 100, and values of -s between 100 and 1000. For seq2HLA, we will determine the effect of adjusting the -3 parameter for values of 0 (default), 5, 10, and 15. For OptiType, we will determine the effect of adjusting the --beta command from values of 0.005 to 0.0015. For HLA-VBseq, we will determine the effect of adjusting the -- alpha_zero parameter between values of 0.01 and 0.10. For HLA-HD, we will determine the effect of adjusting the -c parameter between values of 0.8 and 1.0. For HISAT-genotype, we will determine the effect of adjusting the --simulate-interval parameter for values of 5 and 50, and the read-len parameter for values of 50 and 150. For arcasHLA, we will determine the effect of toggling the --unmapped parameter to True and False. For HLApers, we will determine the effect of toggling the --kallisto option (default star/Salmon) for both creating index, and genotyping. For RNA2HLA, we will determine the effect of adjusting the −3 command for values from 0 to 10, and -c command for values between 0.01 and 0.10. For T1K, we will determine the effect of adjusting the --frac command for values between 0.05 and 0.30, and the --crossGeneRate command for values between 0.01 and 0.10. Altogether, for each combination of parameters and each tool we will evaluate for accuracy on 50 samples from Dataset 1. Using this data we will infer the unique set of optimal set of parameters for each tool.

### Converting the output of the HLA Callers to a universal format

Although some tools are capable of imputation at greater resolutions than two-field resolution, we will consider only the one-field and two-field resolution because two-field resolution is the clinically significant level and our datasets all contain gold standard types with a minimum of two-field resolution. We will first parse all individual output files for each tool to determine the 2 allele imputations at each locus of each sample. For tools that have output ambiguities or more than two allele predictions at a locus, we select the two alleles with highest quality scores. We aggregate all allele imputations for each dataset and convert them into a standard csv format which parallels the format of the gold standard datasets, with each row representing a different sample and each column representing a different loci (see “Determination of Gold Standard HLA types” section of Methods). We will consider lack of prediction as an incorrect prediction. Descriptions of the custom formats for each tool are provided in Table S4. All scripts to convert the tools’ custom results formats to the gold standard format will be made available at https://github.com/Mangul-Lab-USC/HLA_benchmark.

### Evaluating accuracy of HLA predictions

We will evaluate accuracy of each HLA tool in comparison to the gold standard using the following equation:

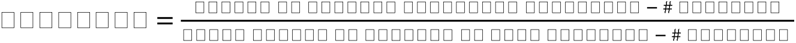

We will assess accuracy at one- and two-field resolutions, with one-field resolution considering only the segment of the HLA allele up to the second field (Figure 1). We will treat no-calls as missed-calls, with the exception of OptiType, which is designed to make only Class I predictions.

Accuracy is determined via a sample-by-sample, locus-by-locus comparison of the prediction with the gold standard. Due to the diploid nature of the loci, the first step is to determine whether to flip the order of the two prediction alleles before comparison with the two gold standard alleles. We will opt to take a “benefit of the doubt” approach, and select the orientation that will maximize the accuracy. To specify, we will perform both a parallel and a crosswise comparison between the allele pair in the gold standard and the prediction, assigning a raw accuracy score to each of the two comparisons. The raw score assigned to each tool considers both one and two-field resolution, weighted to value predictions accurate to two-field resolution over those that are accurate to only one-field resolution. Then, we will select the comparison (either parallel or crosswise) that yields the highest raw score. This orientation of HLA calls to the gold standard will be used for calculation of both the low-resolution and high-resolution accuracy.

We will determine the “# filtered” by counting the number of HLA calls that predict an allele type not present in the gold standard allele list. This “# filtered” value will be subtracted from the numerator and denominator to exclude those alleles from the accuracy calculation.

We will determine the “Number of Alleles Correctly Predicted” by comparing each allele imputation with the corresponding gold standard allele and counting all alleles that represent a match. obtain the numerator to the accuracy function. We will subtract the “# Filtered” from this value to obtain the numerator, and subtract the “# Filtered” from the total number of allele imputations to obtain the denominator. Dividing numerator by denominator will yield the final accuracy. An accuracy of 1 indicates perfect performance, and an accuracy of 0 indicates that every allele is a miscall.

Some gold standard HLA types in Dataset 1 contain allele ambiguities, and in such cases, we consider a match between the prediction with any of the possible alleles to be a fully accurate match. For instance, suppose a loci in the gold standard contained 4 possible alleles, “A*01:01/A*01:04/A*01:22”. An HLA caller that predicted A*01:01 and one that predicted A*01:04 would both be equally and fully accurate.

We plan to examine for statistically significant differences in accuracy across various parameters and features. We will stratify accuracy by features such as class, loci, and allele levels to examine for variations. To assess for statistical significance we will primarily use the chi-square test of independence, with the null hypothesis that there is no difference between levels of each feature examined (class, loci, allele) and the counts of correctly and incorrectly predicted alleles. We also plan to examine the presence of specific alleles that are commonly mispredicted as another specific allele, followed by pairwise alignment of the sequences with the hypothesis that these “miscall pairs” are associated with high sequence similarity.

### Computational modification of RNA-Seq sample properties

We will use Seqtk v1.3 to perform the computational read length shortening on 4 samples of Dataset 1 and 3 samples from Dataset 2, and 4 samples from Dataset 4, for a total of 11 samples (listed in Table S2). Samples from these 3 datasets were chosen because they hace the longest read lengths. We will first convert each sample from FASTQ to FASTA format using the command seqtk seq -a in.fq > out.fa. For each sample, we will then iteratively invoke the command ‘seqtk trimfq -b 5 in.fa > out.fa’, in which each iteration will truncate 5 base pairs from the right of every read, stopping when reads become shorter than 36 bp. Each sample will hence produce multiple computationally modified files, each with read lengths in the range 36bp to original read length with a step size of 5bp. The final files will be converted from fa back to fq format using the seqtk seq -a in.fa > out.fq command. All samples will then be run on the HLA caller tools with default and optimal parameters for accuracy analysis.

### Assessing the effect of ancestral diversity on HLA predictions

We will use all samples in Dataset 1 (n=490) in order to evaluate differences in performances between European and African ancestry groups as dataset 1 is the only dataset for which we have access to the ancestral information. For each caller, we will determine the accuracy at each HLA locus separately for the European subset (n=423) and the African subset (n=67) of dataset 1, both at one- and two-field resolutions. Once the accuracy is calculated, we will use a chi-square test of independence to determine the statistical significance of the differences in accuracy across both ancestral groups. We will also isolate the alleles to investigate the misclassification rate for each allele when looking at European versus African samples.

### Assessing computational resources required to run HLA callers

For each tool, CPU time and RAM metrics were recorded to determine the computational performance for 2 samples in each of Datasets 1-6, for a total of 12 samples (listed in Table S2). Each tool was run with 1 node, 1 task and 2 CPUs per task on the Discovery high-performance computing cluster in the USC Center for Advanced Research Computing (USC CARC). To account for fluctuations in resource availability in the cluster, we averaged the CPU and RAM of each sample across 10 runs. Metrics were collected using an inbuilt command from the cluster’s Slurm Workload Manager, called upon by adding the following line of code, ‘#SBATCH --mail-user=<e-mail address>’ to the slurm batch job script header, would email the user a full record of the execution of the script, containing information on the CPU time and the RAM usage.

## Acknowledgments

We thank Abhinav Nellore, Tianze Tao, Jieting Hu, Ryan Alomair, and Kunali Hepani for their valuable feedback and discussion.

## Data Availability

The RNA-seq datasets 1-7 used for this analysis are available under the following accession numbers on the NCBI SRA archive: ERP001942^35^, SRP298704^36^, ERP000101^37^, SRP099568^38^, SRP198497^39^, SRP096329^40^, and SRP162214^41^. All data required to produce the figures and analysis performed in this paper are freely available at https://github.com/Mangul-Lab-USC/HLA_benchmark. We are committed to sharing all raw data and materials collected in-house upon acceptance of Stage 2 manuscript.

## Code Availability

All code required to produce the figures and analysis performed in this paper is under the Massachusetts Institute of Technology (MIT) license and will be freely available at https://github.com/Mangul-Lab-USC/HLA_benchmark. We are committed to sharing all code on acceptance of your Stage 2 manuscript, including code used to conduct power analyses and analyze pilot data should be made accessible in the same location.

## Supplementary

### Supplementary Figures

**Figure S1:**
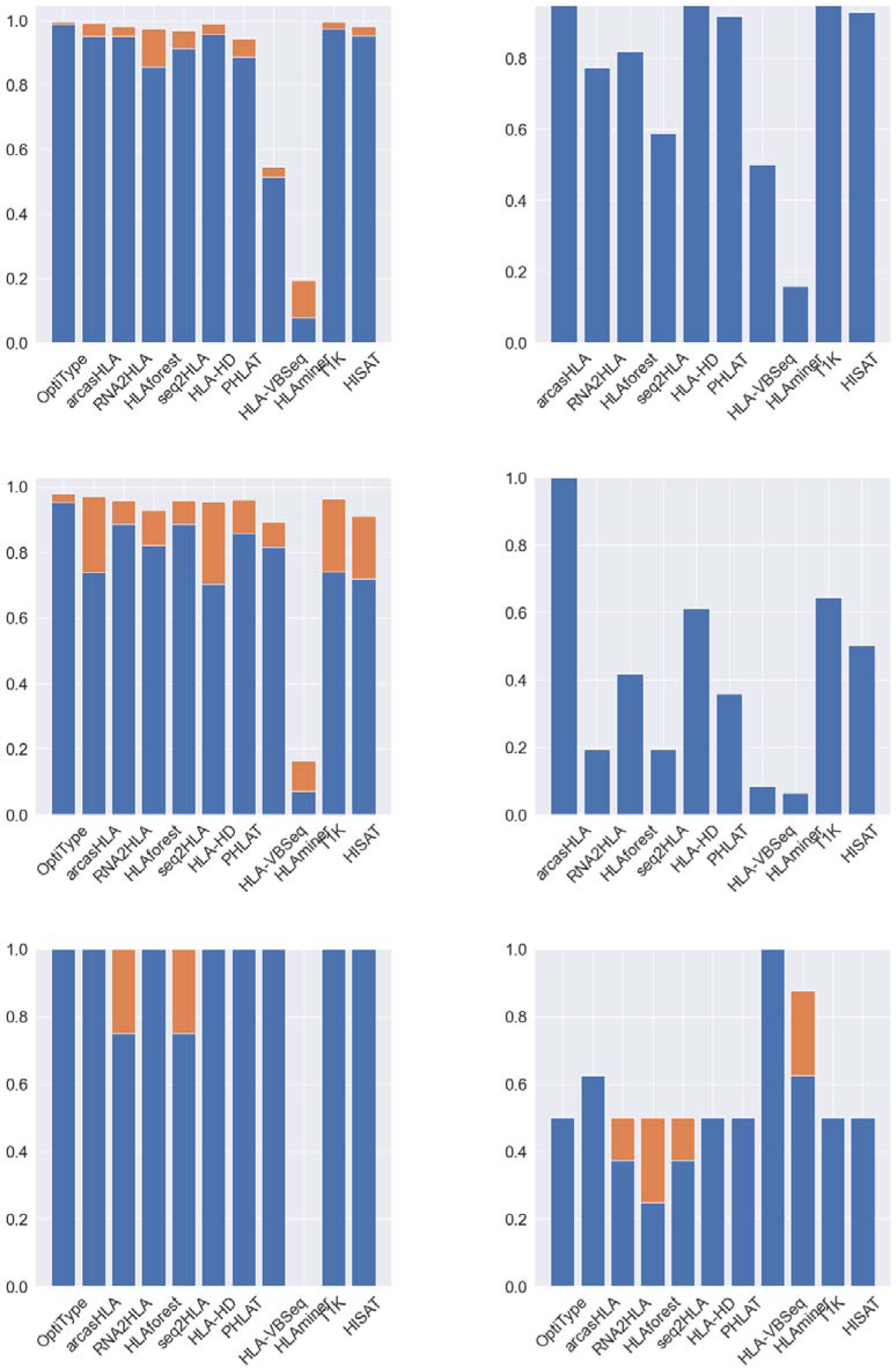
accuracy for each dataset individually

**Figure S2:**
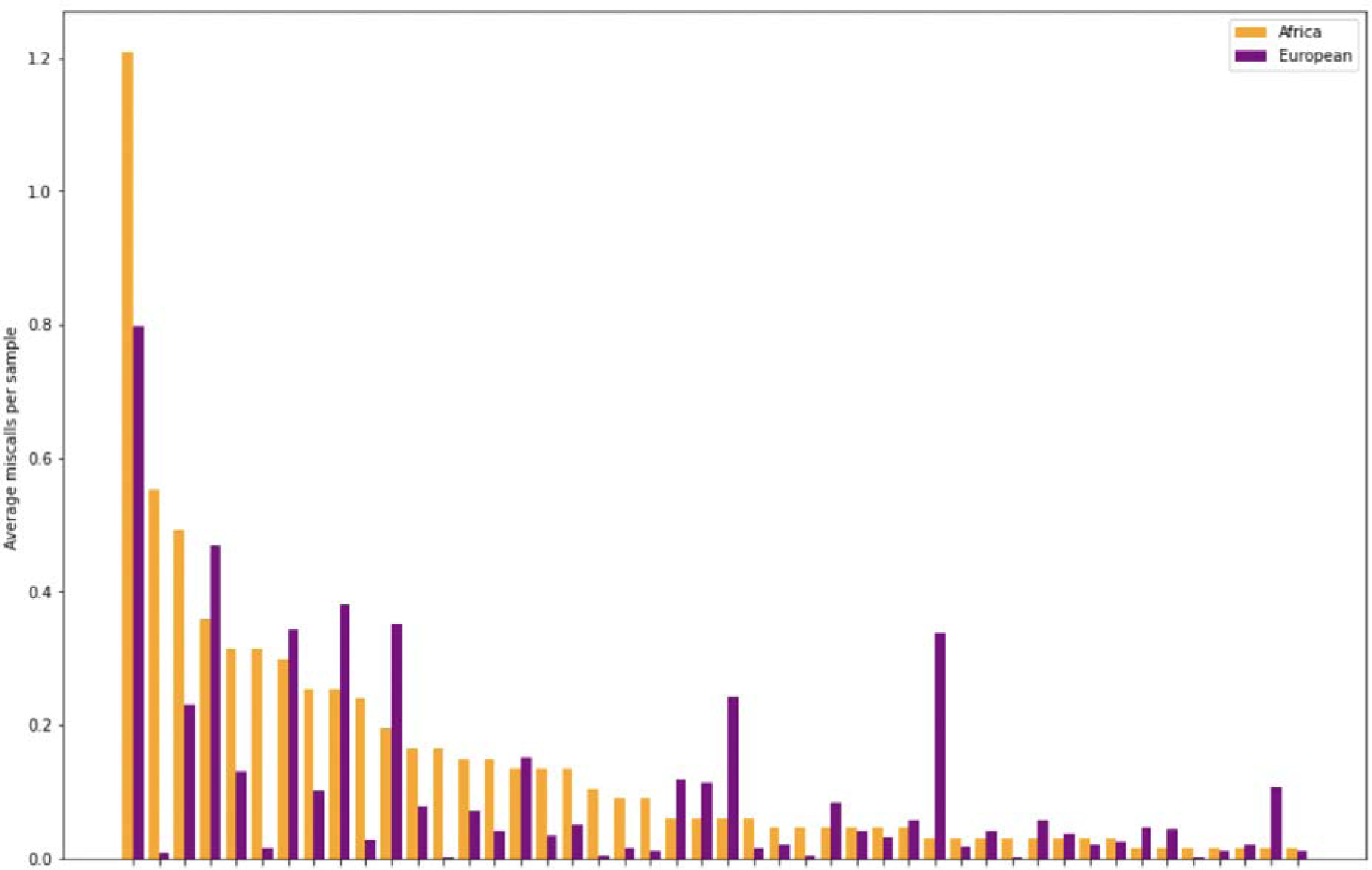
% miscalls for each allele, for Africa vs Europe ancestry

### Supplementary Tables

**Table S1:**
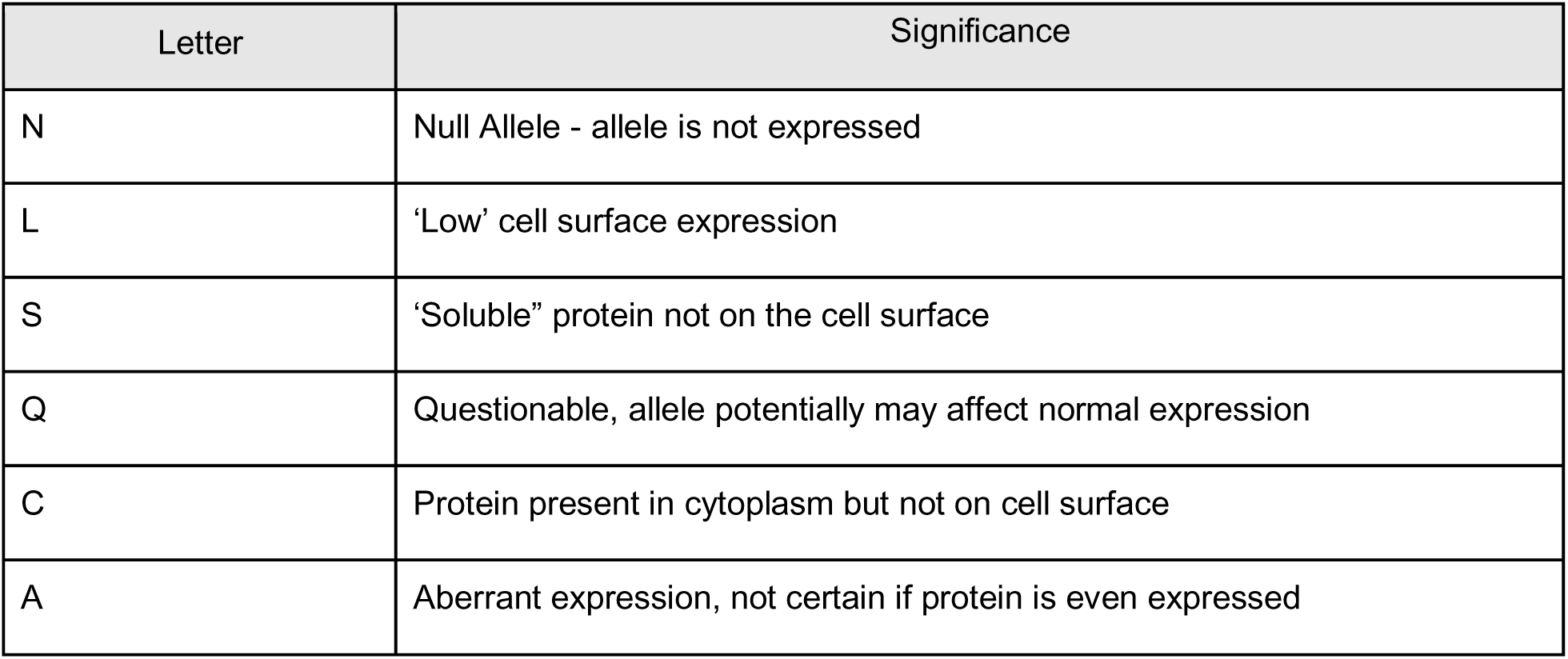
The Suffix Letter nomenclature for the specification of the different types of expression of HLA alleles. Typically the Suffix Letter is at the end of the HLA Name, but can be included at the two-field resolution.

**Table S2:**
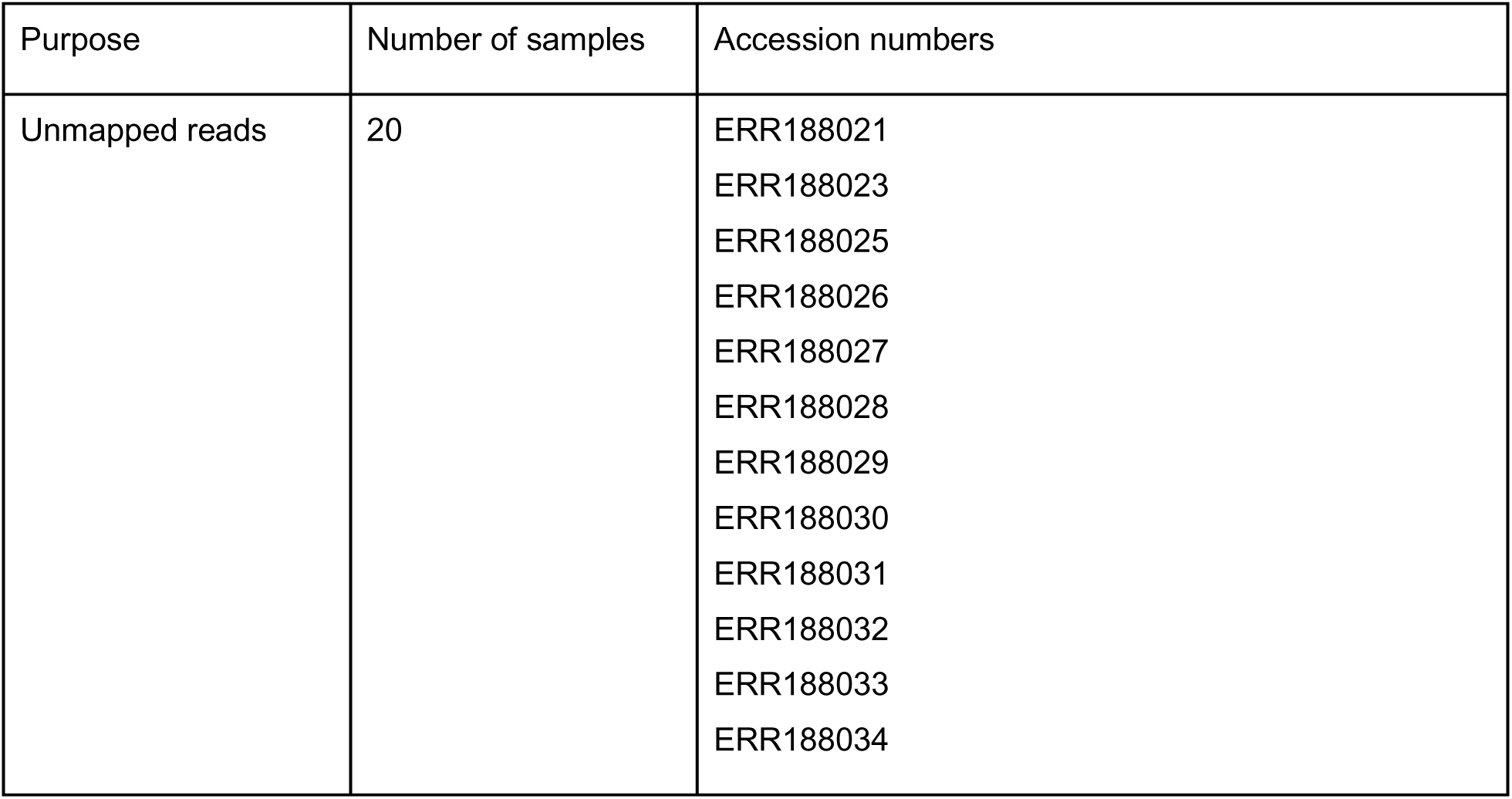
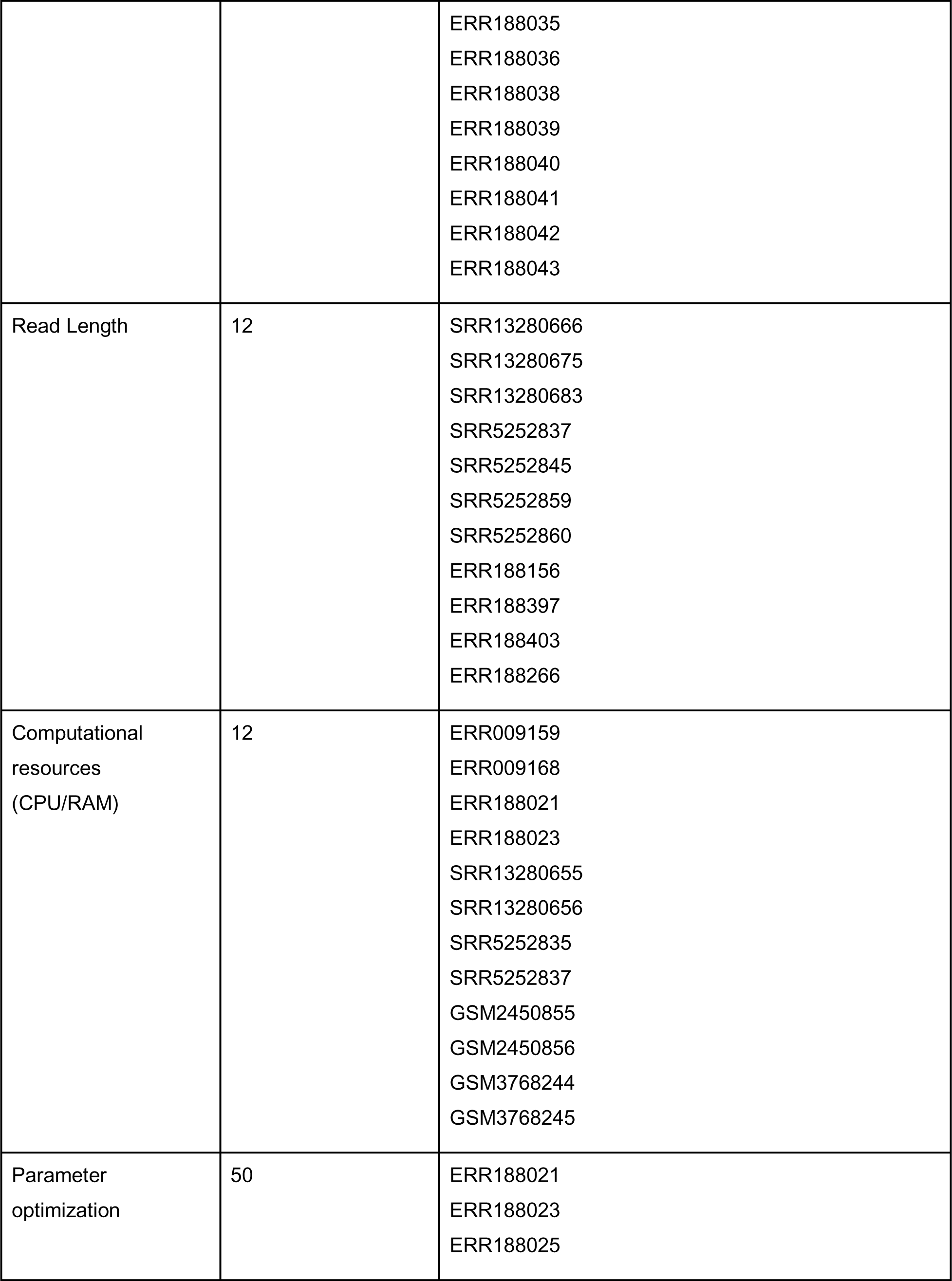
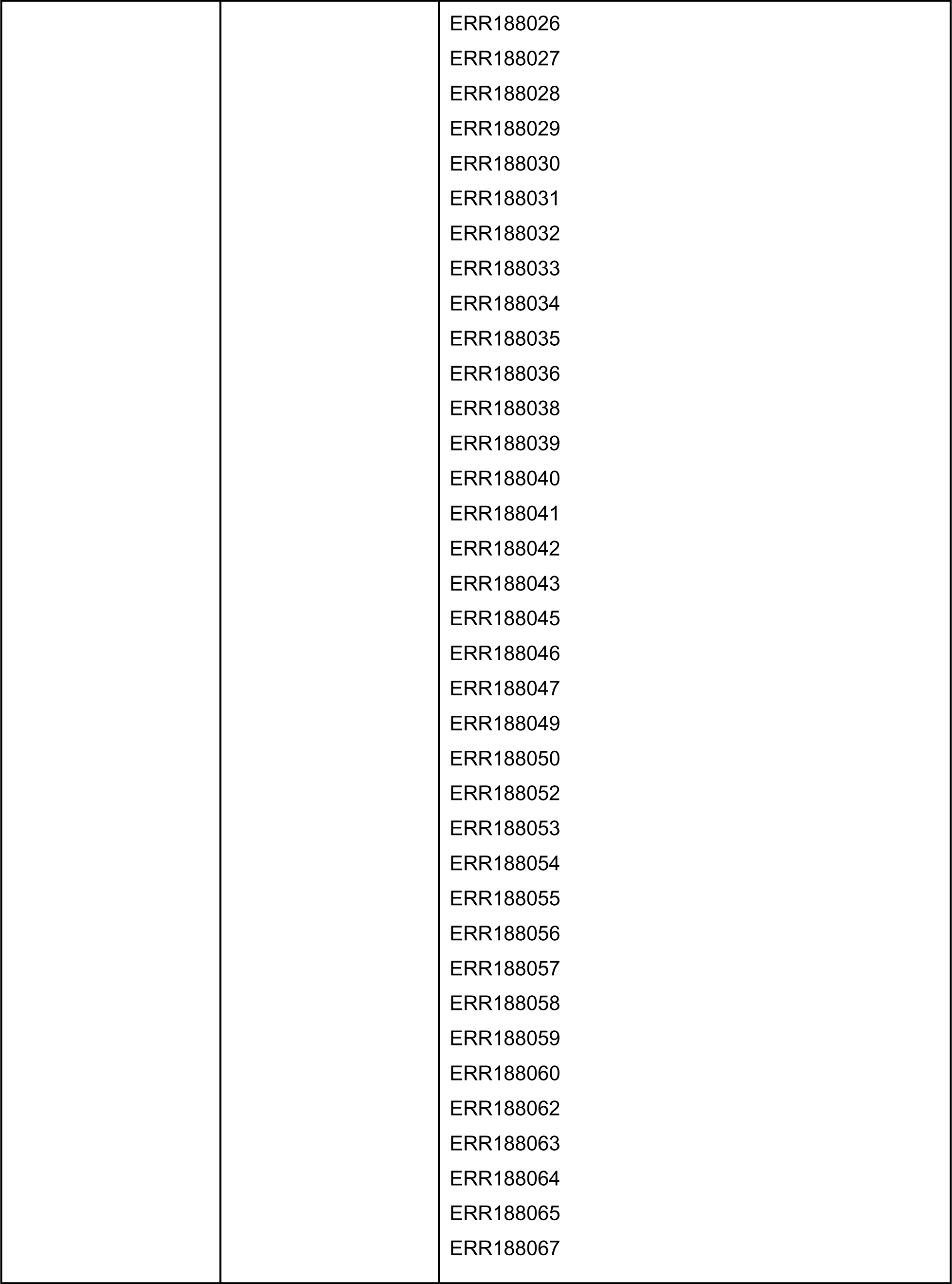
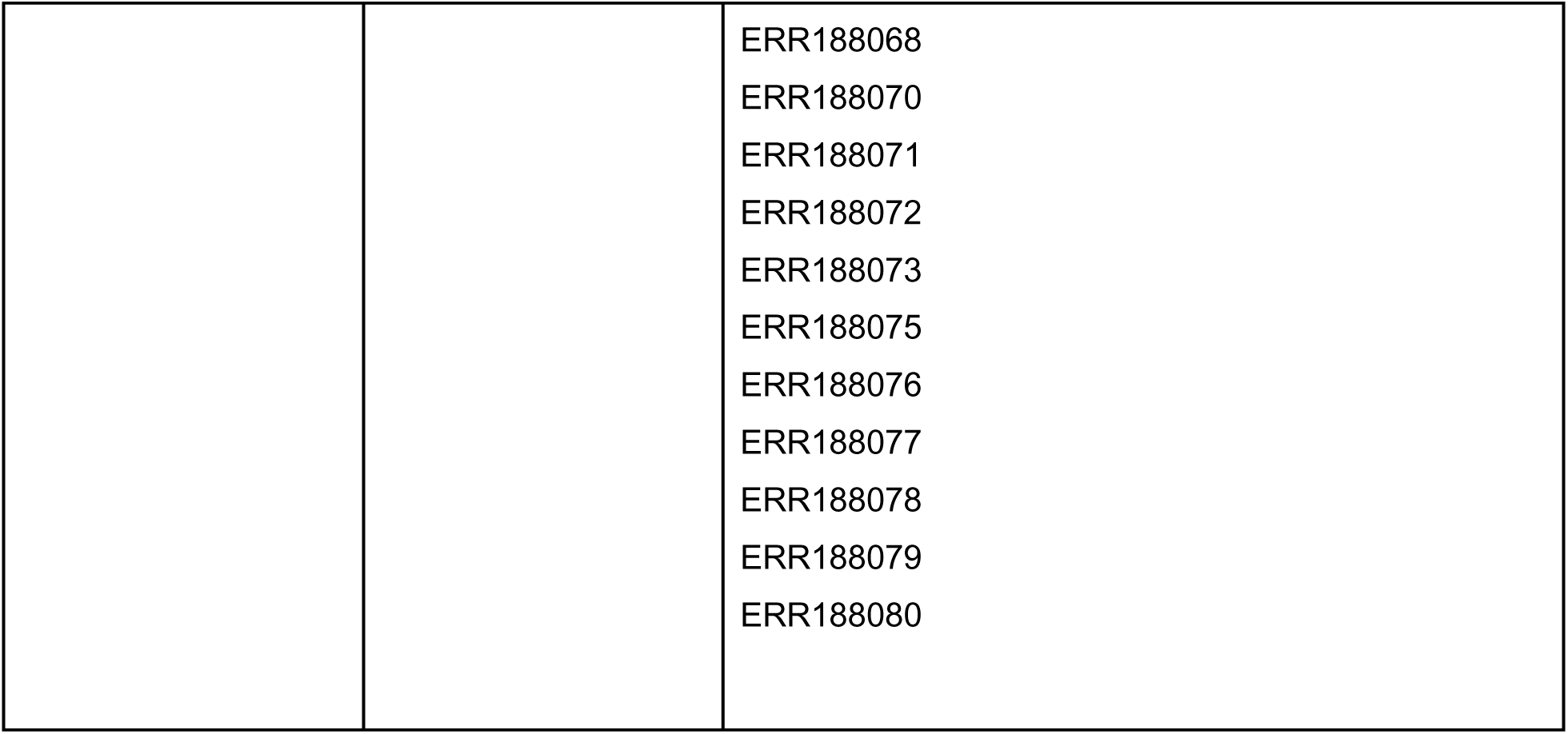
Accession numbers to subsampled datasets used for some accuracy benchmarks. “Purpose” lists the benchmarking purpose for the subsampled dataset. “Number of samples” denotes the number of samples in the subsampled dataset, and “Accession numbers” lists the specific SRA accessions to those samples.

**Table S3:**
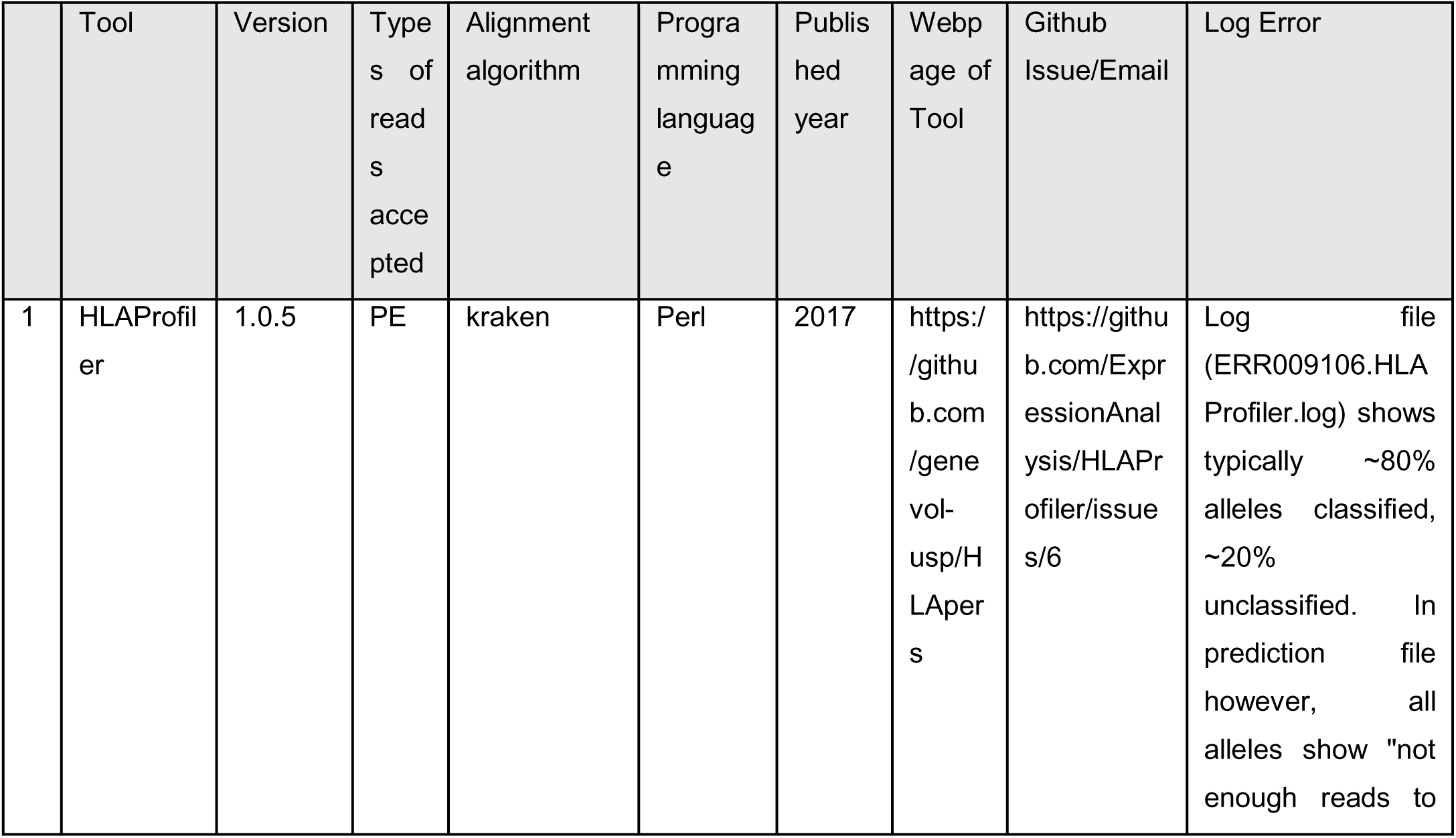
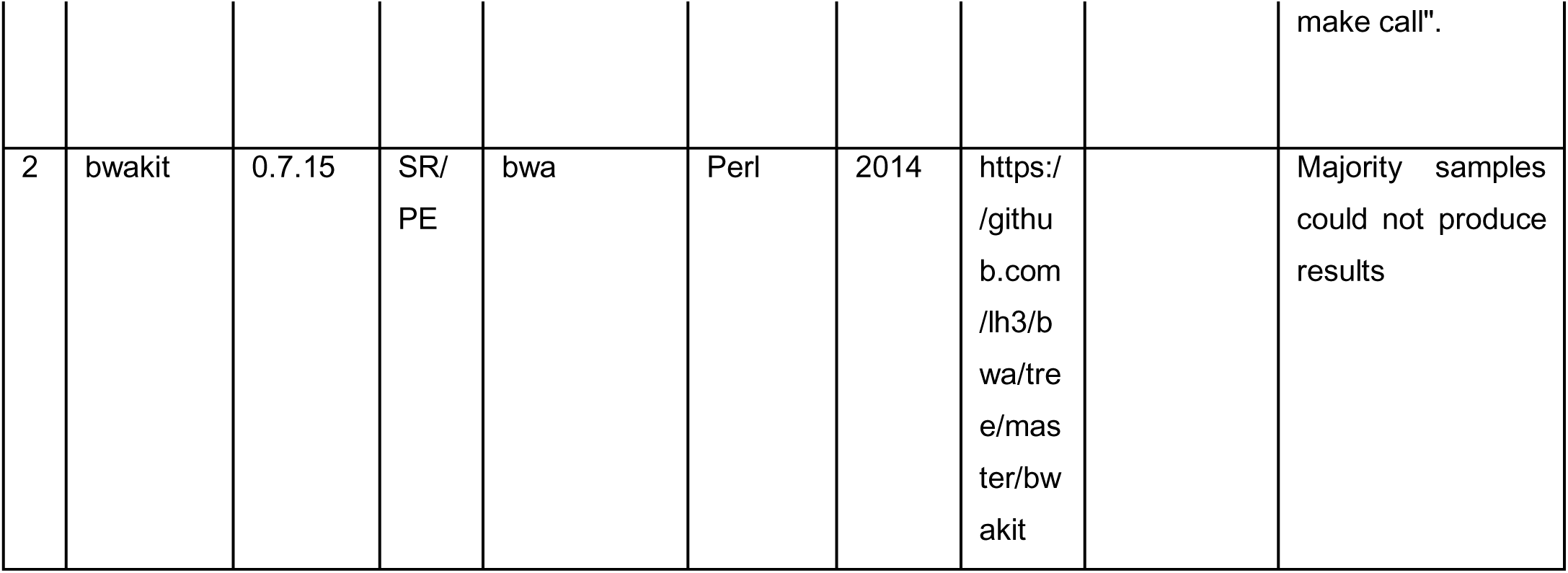
Existing Methods for HLA typing which we failed to install or run. We opened an issue on each tool’s GitHub if available. If no GitHub was publicly available, then we emailed the corresponding authors of the paper for each tool.

**Table S4:**
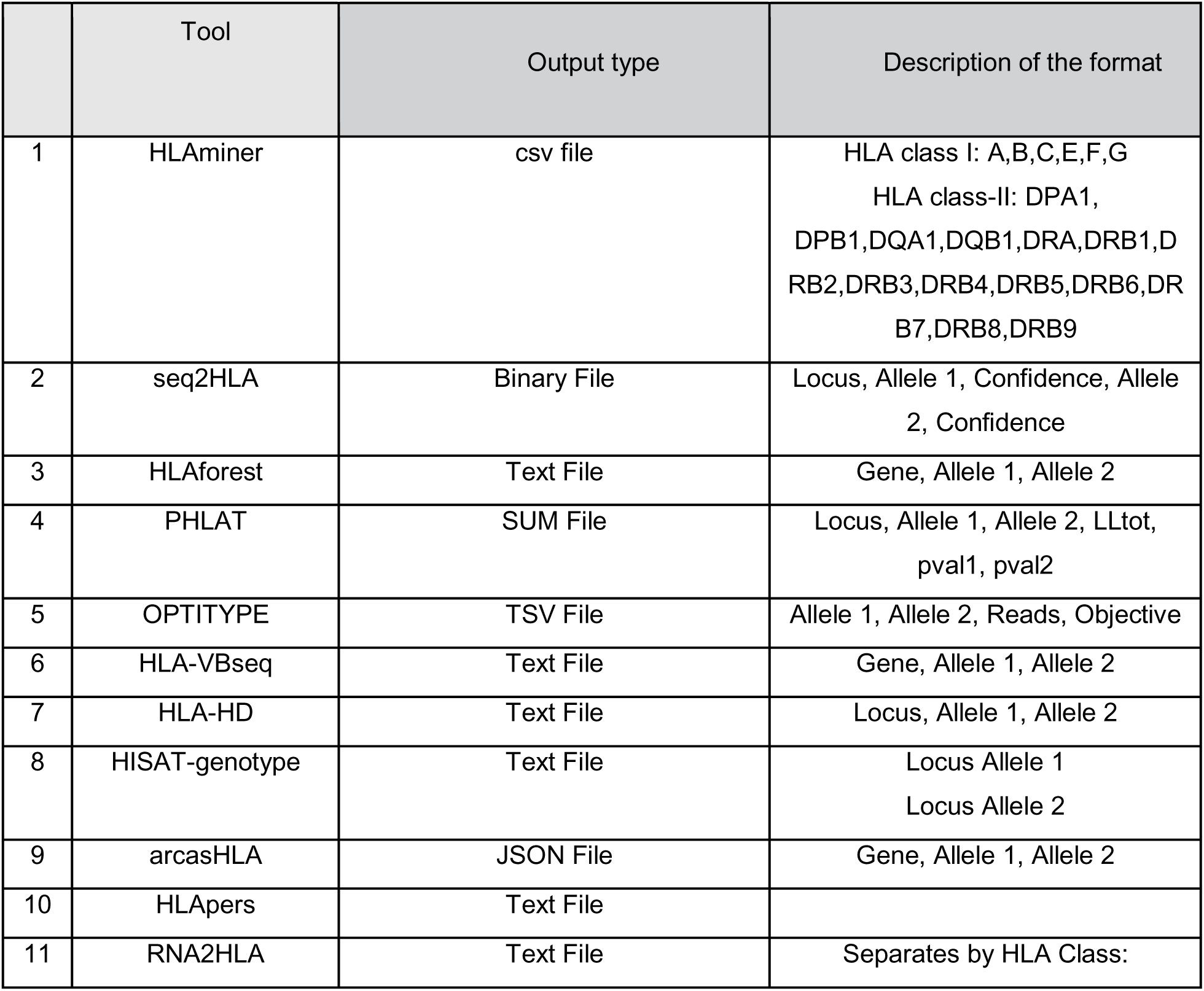
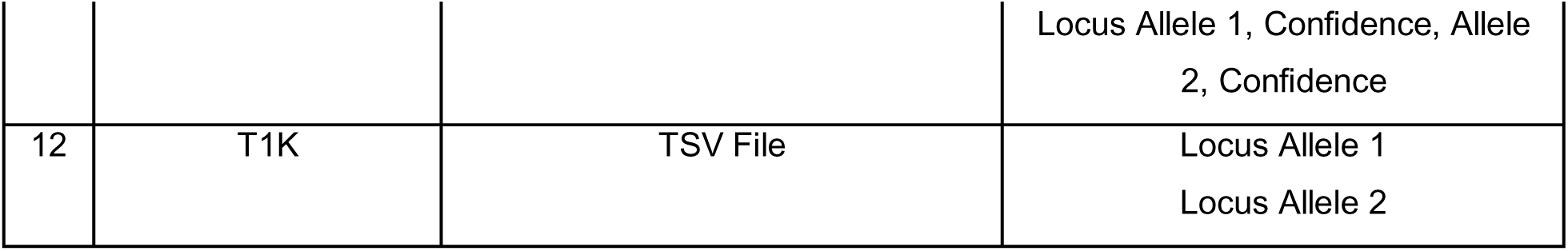
The Raw Output Types of each caller and a description of the output organization within the file.

**Table S5:** accuracy for (a) class I (b) class II (c) overall.

| Tool | % inaccurate | % one-field | % two-field |
| --- | --- | --- | --- |
| **OptiType** | 0 | 0 | 0 |
| **arcasHLA** | 0.016144 | 0.049383 | 0.934473 |
| **RNA2HLA** | 0.067753 | 0.048885 | 0.883362 |
| **HLAforest** | 0.024332 | 0.11021 | 0.865458 |
| **seq2HLA** | 0.085835 | 0.094278 | 0.819887 |
| **HLA-HD** | 0.062383 | 0.030019 | 0.907598 |
| **PHLAT** | 0.109276 | 0.064168 | 0.826557 |
| **HLA-VBseq** | 0.926346 | 0.013692 | 0.059962 |
| **HLAminer** | 0.763615 | 0.236385 | 0 |

B
| Tool | % inaccurate | % one-field | % two-field |
| --- | --- | --- | --- |
| **OptiType** | 0.006173 | 0.011111 | 0.982716 |
| **arcasHLA** | 0.014763 | 0.050385 | 0.934852 |
| **RNA2HLA** | 0.015625 | 0.032813 | 0.951562 |
| **HLAforest** | 0.036814 | 0.124163 | 0.839023 |
| **seq2HLA** | 0.015625 | 0.032813 | 0.951562 |
| **HLA-HD** | 0.058768 | 0.046512 | 0.89472 |
| **PHLAT** | 0.027177 | 0.061231 | 0.911591 |
| **HLA-VBseq** | 0.099871 | 0.05139 | 0.848739 |
| **HLAminer** | 0.850141 | 0.085684 | 0.064175 |

C
| Tool | % inaccurate | % one-field | % two-field |
| --- | --- | --- | --- |
| **OptiType** | 0.006173 | 0.011111 | 0.982716 |
| **arcasHLA** | 0.01532 | 0.049981 | 0.934699 |
| **RNA2HLA** | 0.029546 | 0.037105 | 0.933349 |
| **HLAforest** | 0.031668 | 0.118411 | 0.849921 |
| **seq2HLA** | 0.043698 | 0.057389 | 0.898912 |
| **HLA-HD** | 0.060218 | 0.039895 | 0.899887 |
| **PHLAT** | 0.055099 | 0.06223 | 0.882671 |
| **HLA-VBseq** | 0.435725 | 0.036071 | 0.528204 |
| **HLAminer** | 0.817405 | 0.142701 | 0.039895 |

**Table S6a:**
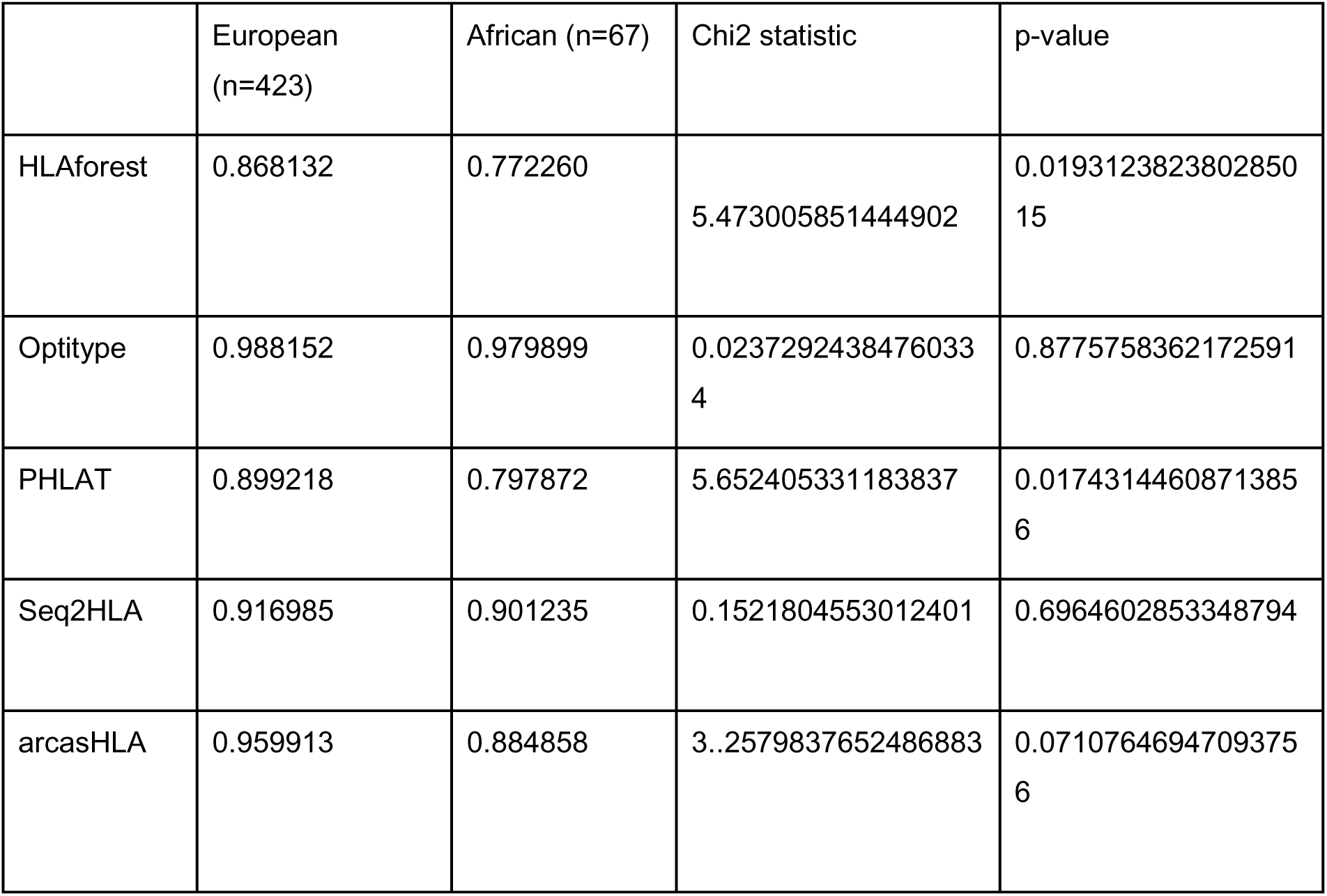
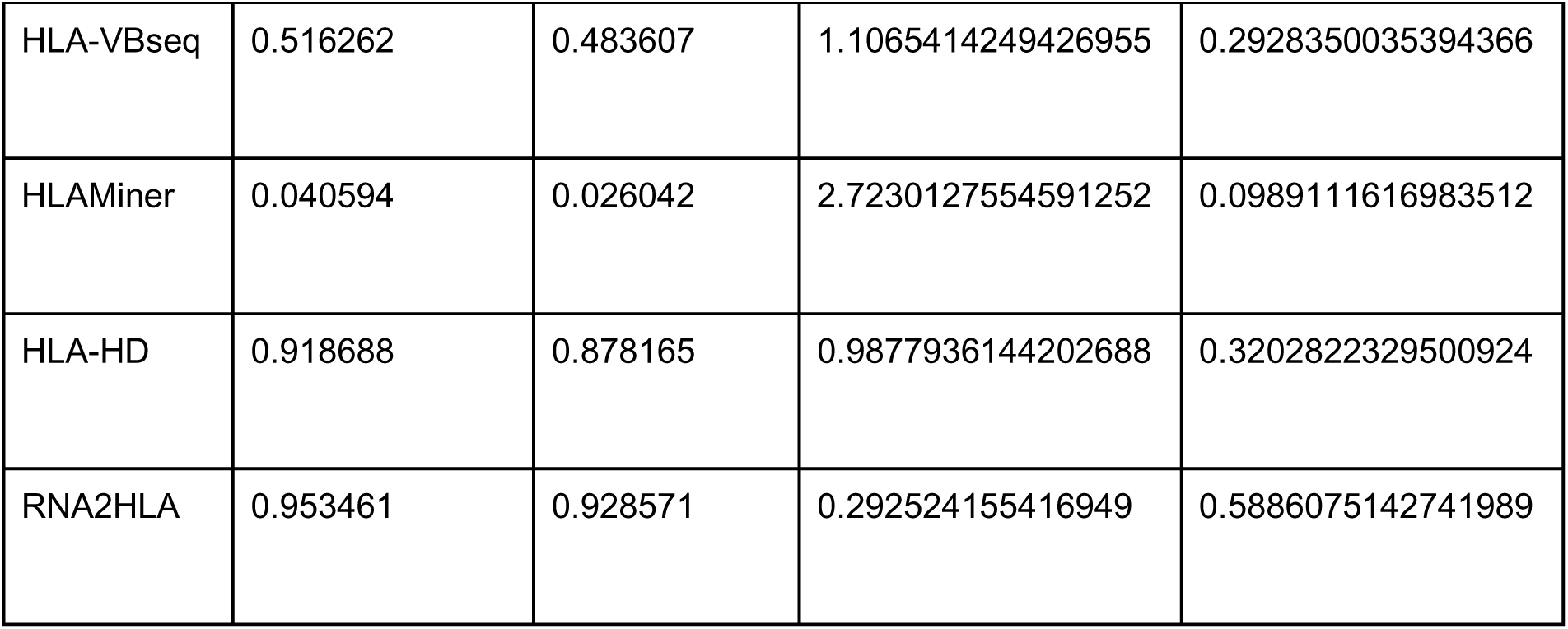
Two-field Accuracy of HLA callers, for European and African ancestry, to two-field significance (individual tools is chi square test of independence)

**Table S6b:**
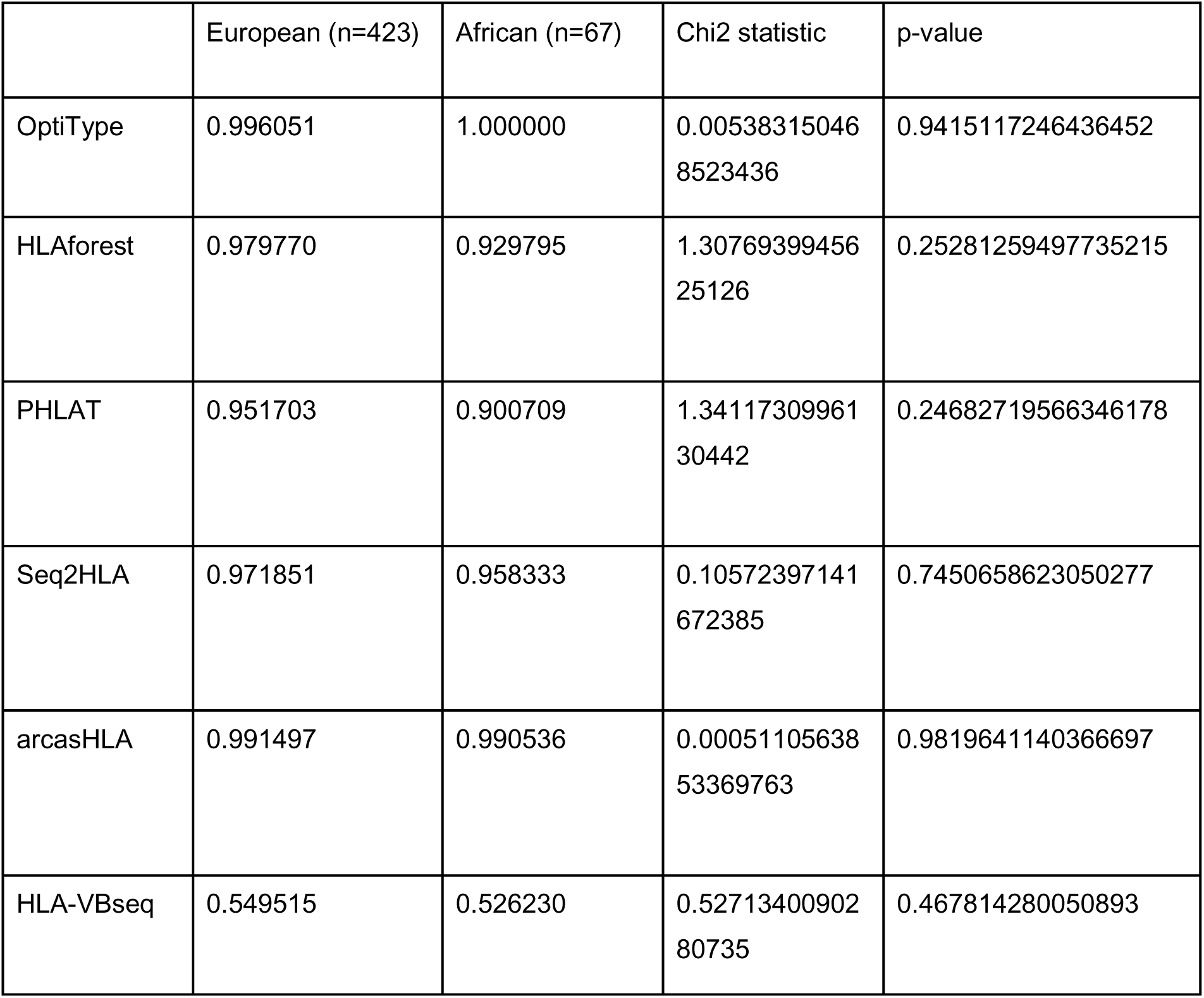
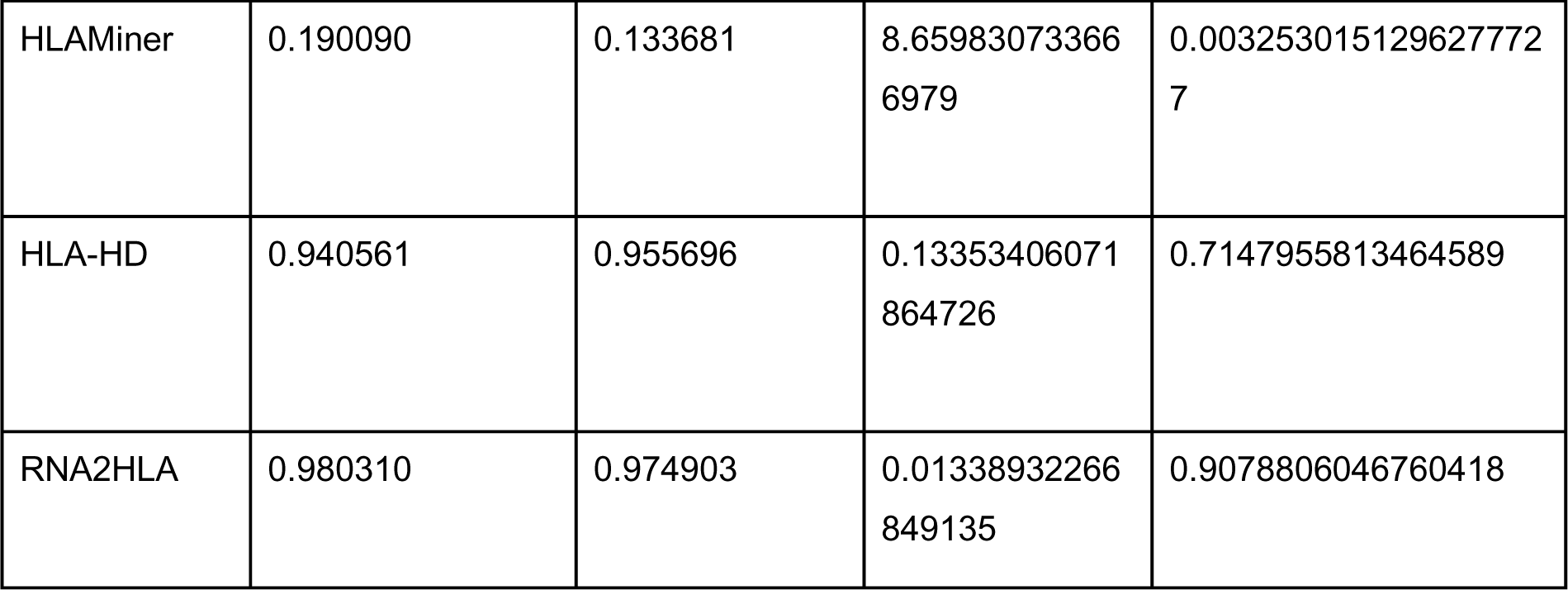
Two-field Accuracy of HLA callers, for European and African ancestry, to one-field significance (overall is binomial test, individual tools is chi square test of independence)

|  | European (n=423) | African (n=67) | Chi2 statistic | p-value |
| --- | --- | --- | --- | --- |
| OptiType | 0.996051 | 1.000000 | 0.00538315046<br>8523436 | 0.9415117246436452 |
| HLAforest | 0.979770 | 0.929795 | 1.30769399456<br>25126 | 0.25281259497735215 |
| PHLAT | 0.951703 | 0.900709 | 1.34117309961<br>30442 | 0.24682719566346178 |
| Seq2HLA | 0.971851 | 0.958333 | 0.10572397141<br>672385 | 0.7450658623050277 |
| arcasHLA | 0.991497 | 0.990536 | 0.00051105638<br>53369763 | 0.9819641140366697 |
| HLA-VBseq | 0.549515 | 0.526230 | 0.52713400902<br>80735 | 0.467814280050893 |
| HLAMiner | 0.190090 | 0.133681 | 8.65983073366<br>6979 | 0.003253015129627772<br>7 |
| HLA-HD | 0.940561 | 0.955696 | 0.13353406071<br>864726 | 0.7147955813464589 |
| RNA2HLA | 0.980310 | 0.974903 | 0.01338932266<br>849135 | 0.9078806046760418 |

